# Exifone is a Potent HDAC1 Activator with Neuroprotective Activity in Human Neuronal Models of Neurodegeneration

**DOI:** 10.1101/2020.03.02.973636

**Authors:** Debasis Patnaik, Ping-Chieh Pao, Wen-Ning Zhao, M. Catarina Silva, Norma K. Hylton, Peter S. Chindavong, Ling Pan, Li-Huei Tsai, Stephen J. Haggarty

**Author notes:** Corresponding Author, Correspondence should be addressed to: Stephen J. Haggarty, Associate Professor of Neurology, Massachusetts General Hospital, Harvard Medical School, Center for Genomic Medicine, 185 Cambridge Street, Boston, MA, 02114, USA.

## Abstract

Genomic instability caused by a deficiency in the DNA damage response and repair has been linked to age-related cognitive decline and neurodegenerative diseases. Preventing genomic instability that ultimately leads to neuronal death may provide a broadly effective strategy to protect against multiple potential genotoxic stressors. Recently, the zinc-dependent, class I histone deacetylase HDAC1 has been identified as a critical factor for protecting neurons from deleterious effects of DNA damage in Alzheimer’s disease (AD), amyotrophic lateral sclerosis (ALS), and frontotemporal dementia (FTD). Translating these observations to a novel neuroprotective therapy for AD, ALS, and FTD may be advanced by the identification of small molecules capable of increasing the deacetylase activity of HDAC1 selectively over other structurally similar HDACs. Here, we demonstrate that exifone, a drug previously shown to be effective in treating cognitive deficits associated with AD and Parkinson’s disease, the molecular mechanism of which has remained poorly understood, potently activates the deacetylase activity of HDAC1 and provides protection against genotoxic stress. We show that exifone acts as a mixed, non-essential activator of HDAC1 that is capable of binding to both free and substrate-bound enzyme resulting in an increased relative maximal rate of HDAC1-catalyzed deacetylation. Exifone can directly bind to HDAC1 based upon biolayer interferometry assays with kinetic and selectivity profiling suggesting HDAC1 is preferentially targeted compared to other class I HDACs and the kinase CDK5 that have also been implicated in neurodegeneration. Consistent with a mechanism of deacetylase activation intracellularly, treatment of human induced pluripotent stem cell (iPSC)-derived neuronal cells resulted in globally decreased histone acetylation. Moreover, exifone treatment was neuroprotective in a tauopathy patient iPSC-derived neuronal model subject to oxidative stress. Taken together, these findings reveal exifone as a potent activator of HDAC1-mediated deacetylation, thereby offering a lead for novel therapeutic development aiming to protect genomic integrity in the context of neurodegeneration and aging.

**Graphical Abstract:** 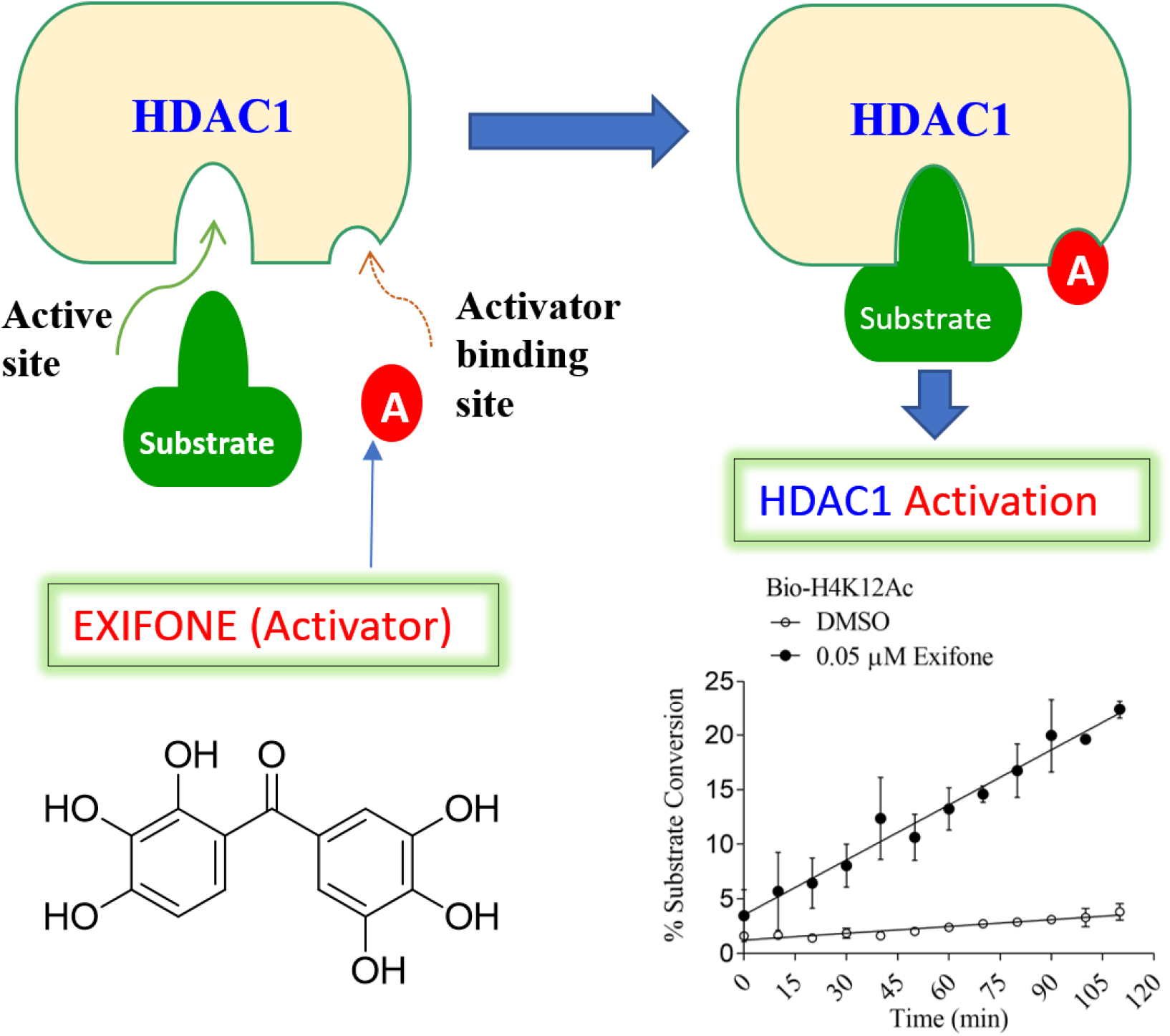

## Introduction

Alzheimer’s disease (AD) and cognitive disorders are common in the elderly population and are frequently caused by neurodegeneration that leads to a progressive and currently irreversible decline of plasticity and function of the central nervous systems (CNS). Medications approved by the Food and Drug Administration (FDA) for the treatment of AD are largely symptomatic in nature and include cholinesterase inhibitors such as donepezil (Aricept), rivastigmine (Exelon), galantamine (Razadyne); and memantine (Namenda), an NMDA (*N-methyl-D-aspartate*) receptor antagonist. None of these medications prevent disease progression necessitating the discovery of effective disease-modifying agents with a unique mechanism of action.

Activation of histone deacetylase 1 (HDAC1) has emerged as a new strategy to prevent neurodegeneration by blocking cell cycle re-entry and preventing DNA damage in the context of neurodegeneration, including AD^1–2^ and amyotrophic lateral sclerosis^3^. Whereas overexpression of catalytically active HDAC1, but not a mutant lacking deacetylase activity, conferred protection against p25/CDK5-mediated DNA damage and neuronal death in a mouse model of stroke^1^. Moreover, inhibition of HDAC1 catalytic activity has been observed in an inducible p25/CDK5 neurodegeneration mouse model, which exhibits critical pathological signatures of AD, including loss of neurons in the forebrain, memory impairment, and increased β-amyloid peptide production. Togethe^r1-3^ these findings indicate a role for HDAC1 in the maintenance of genomic integrity and neuroprotection.

To advance the therapeutic application of these observations concerning the neuroprotective role of HDAC1, we have identified multiple chemical classes of small molecule activators of HDAC1 via high throughput screening^4^. Our recent characterization of HDAC1 small-molecule activators from a primary screen using *in vitro* enzymatic activity and structure-activity relationship (SAR) studies led us to investigate exifone as an HDAC1 activator^4^. Exifone (2,3,3’,4,4’,5’-hexahydroxybenzophenone; Adlone) is a member of the benzophenone class of molecules synthesized in the 1970’s in France that was initially found to have benefit for treatment of blood microcirculation disorders although its precise mechanism of action was not well understood^5^. Exifone was subsequently shown to have neuroprotective activity in a battery of preclinical models and through controlled clinical trials with oral doses ranging from 200 – 600 mg/day shown to be an effective treatment for cognitive deficits in elderly patients with both Alzheimer-type dementia^6^ and Parkinson’s disease leading to its approval as a medication in France in 1988^7^. Additionally, Porsolt and colleagues provided evidence of exifone’s facilitation of memory function in mouse models that was interpreted to be consistent with its proposed effects in humans^8–9^, and additional preclinical studies suggested exifone has properties of an atypical antidepressant. While in these early preclinical and clinical studies exifone was well tolerated and did not produce any noticed sign of toxicity, continued administration of high doses of exifone (i.e., 600 mg/day for 2 to 4 months) resulted in reversible liver damage in about 1 in 15,000 patients leading to its registration being withdrawn in 1990^10–14^.

Exifone has also been shown to have free radical scavenging properties^15^, which led to the synthesis of structurally related derivatives to be investigated as neuroprotective antioxidant molecules^16^. Exifone’s pharmacological mode of action has been reported to include activation of oxygen and glucose metabolism in neurons and antagonism of aminergic neurotransmission impairment induced by temporary ischemia^7, 17^. In neuronal-like PC12 cells, exifone has been shown to reduce^7^ the association of toxic peptides with the cell membranes and to prevent toxicity by β-amyloid peptides^18^.

In this report, we demonstrate for the first time with recombinant enzyme preparations *in vitro* that exifone is a potent activator of HDAC1 with a non-essential mechanism of activation. We further show that exifone treatment of human induced pluripotent stem cell (iPSC)-derived neuronal cells leads to a measurable decrease in histone acetylation. Finally, we demonstrate that exifone treatment can provide neuroprotection in the context of human neurons derived from patient with tauopathy. In a companion study, we show that exifone is able to reduce DNA damage *in vivo* in an AD mouse model and further linking fundamental mechanisms of DNA damage response pathway and repair to HDAC1 activity^2^.

## Results

### Exifone is a potent small molecule activator of HDAC1

Exifone (**Figure 1A**) and its analog benzophenone (**Figure 1B**) that lacks any hydroxyl substituents were tested for their influence on the enzymatic activity of HDAC1. We used a quantitative, mass spectrometry-based assay (RapidFire MS) to measure the deacetylation activity on an acetylated peptide substrate to understand the influence of exifone on the rate of HDAC1 deacetylation. We have found this assay platform has the advantages of allowing direct, and highly sensitive measurement of deacetylated product formation at low substrate concentrations. Moreover, it is not dependent upon using non-physiological, fluorophore-modified peptides and coupling to trypsin for a readout that historically has created challenges when characterizing other classes of deacetylase activators as recently reviewed^19–20^. Using this RapidFire MS platform, measurement of dose-response for exifone effect on HDAC1 activation, using an acetylated peptide derived from histone H4 (Bio-H4K12Ac) and a non-histone substrate (Bio-p53K382Ac), revealed EC_50_ values of 0.045 μM and 0.065 μM, respectively (**Figure 1C and 1E**). Moreover, the top fitted values for deacetylase activity (substrate conversion) were comparatively higher for the Bio-H4K12Ac substrate. Benzophenone did not act as an HDAC1 activator for either of the peptide substrates tested over the concentration range tested, with the resulting HDAC1 activity remaining near the basal level (**Figure 1D and 1F**).

**Figure 1.**
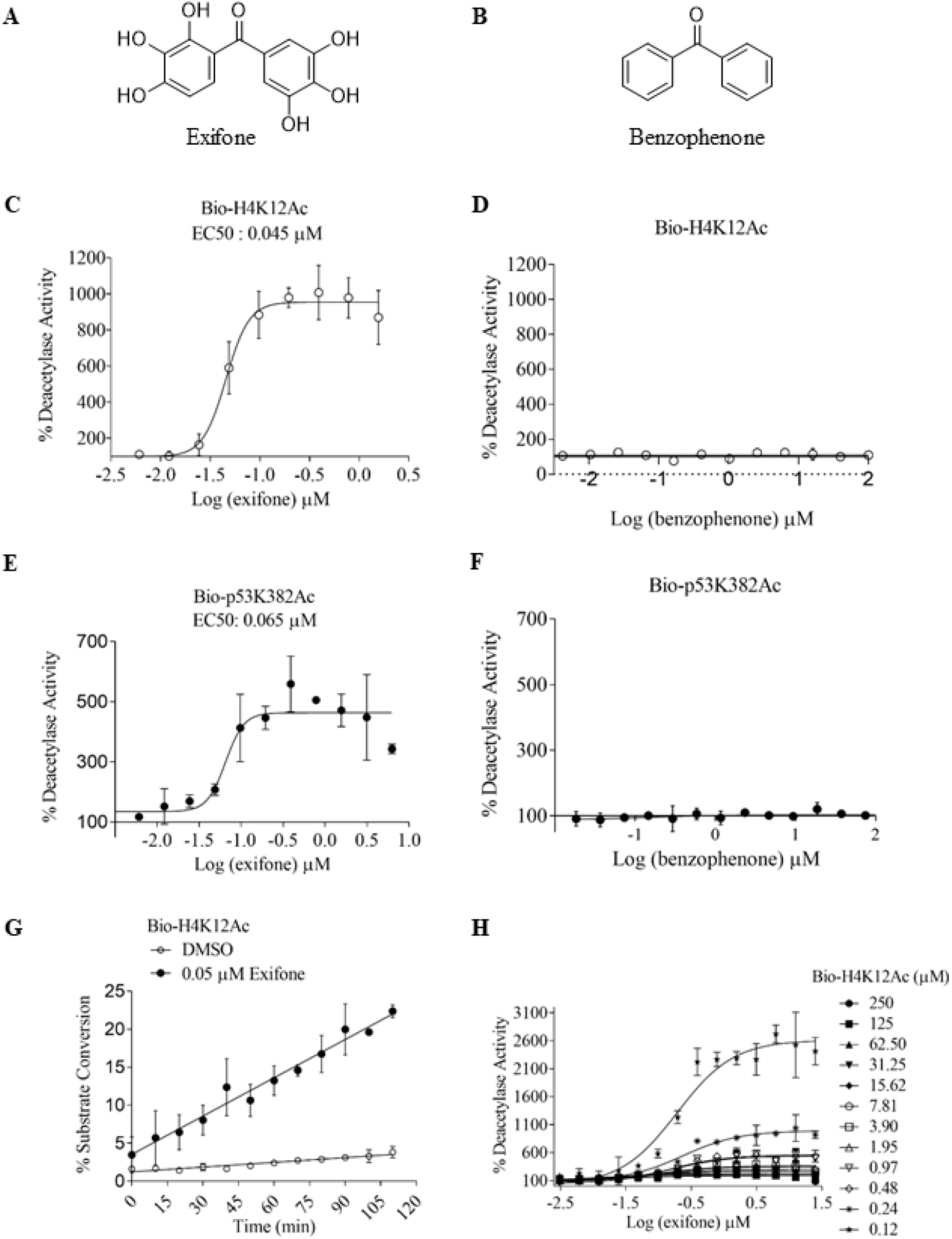
Exifone as a potent small molecule activator of HDAC1. Structure of exifone (**A**) and benzophenone (**B**), respectively. Dose-dependent activation of HDAC1 by exifone in RapidFire mass spectrometry assay for HDAC1 with 1 μM Bio-H4K12Ac substrate (**C**) or 1 μM Bio-p53K382Ac substrate (**E**). The compound benzophenone (**B**), a structurally simpler compound that lacks the six hydroxyl groups does not cause HDAC1 activation (**D** and **F**). **(G)** Exifone increased the rate of deacetylation reaction by HDAC1 as observed via monitoring the percentage of substrate conversion. Mass spectrometry-based (Rapid-Fire) HDAC1 deacetylase activity assay was performed with 1 μM Bio-H4K12Ac substrate and 40 nM enzyme. The figure demonstrates the linearity of reaction progression as a function of time in the absence (control) and the presence of exifone. **(H)** Dose-dependent activation of HDAC1 by exifone at variable concentrations of the acetylated substrate peptide (Bio-H4K12Ac). The highest level of enzyme activation was observed when the substrate concentration is significantly lower than the apparent *K_m_* value for the acetylated peptide substrate ([S]<<[K_m_]). No HDAC1 activation was observed when acetylated substrate peptides were used at higher concentrations such as 125 and 250 μM.>

**Figure 1G** shows the linearity of HDAC1 reaction progression as a function of time in the absence (control) and the presence of exifone. The increase in substrate conversion demonstrates the catalytic activation of HDAC1. Exifone acts as a potent small molecule activator of HDAC1 and thus increased the rate of deacetylation by the enzyme.

### Exifone increases HDAC1 catalytic activity via non-essential activation

The mass spectrometry-based (Rapid-Fire) HDAC1 activity assay was performed with 1 μM of Bio-H4K12Ac substrate and 40 nM of enzyme. Dose-dependent activation of HDAC1 by exifone was determined at various concentrations of the Bio-H4K12Ac substrate. The highest level of HDAC1 activation was observed when the substrate concentration was significantly lower than its *K*_m_ value ([S] << [*K*_m_]). The increase of histone deacetylase activity was more pronounced when the peptide substrate concentration was significantly below the *K*_m_ value (Supplemental Table 1). In contrast, no HDAC1 activation was observed with exifone when peptide substrate concentrations of 125 μM and 250 μM were used (**Figure 1H**).

The reaction mechanism of HDAC1 activation by exifone was determined by simultaneously varying the concentration of both the activator and acetylated substrate. Data were analyzed by nonlinear regression and fitted with the Michaelis-Menten equation to determine *K*_m_ of the acetylated substrate at various concentrations of the activator exifone (**Figure 2A**). Increasing concentrations of exifone caused a progressive decrease of substrate *K*_m_ value, thus demonstrating the role of an activator that increases the affinity of HDAC1 for the Bio-H4K12Ac substrate (**Figure 2B**). Increasing exifone concentration caused a decrease of the apparent substrate *K*_m_ for Bio-H4K12Ac peptide and increased the maximal reaction velocity. Exifone thus appears to increase the affinity of the enzyme for the acetylated substrate peptide, as evidenced by a decrease in substrate *K*_m_ value. The value of alpha (α) and beta (β) were estimated as 0.3 and 1.46, respectively. Alpha is the factor by which substrate *K*_m_ changes when activator interacts with the enzyme, and beta is the factor that reflects *V*_max_ change.

**Figure 2.**
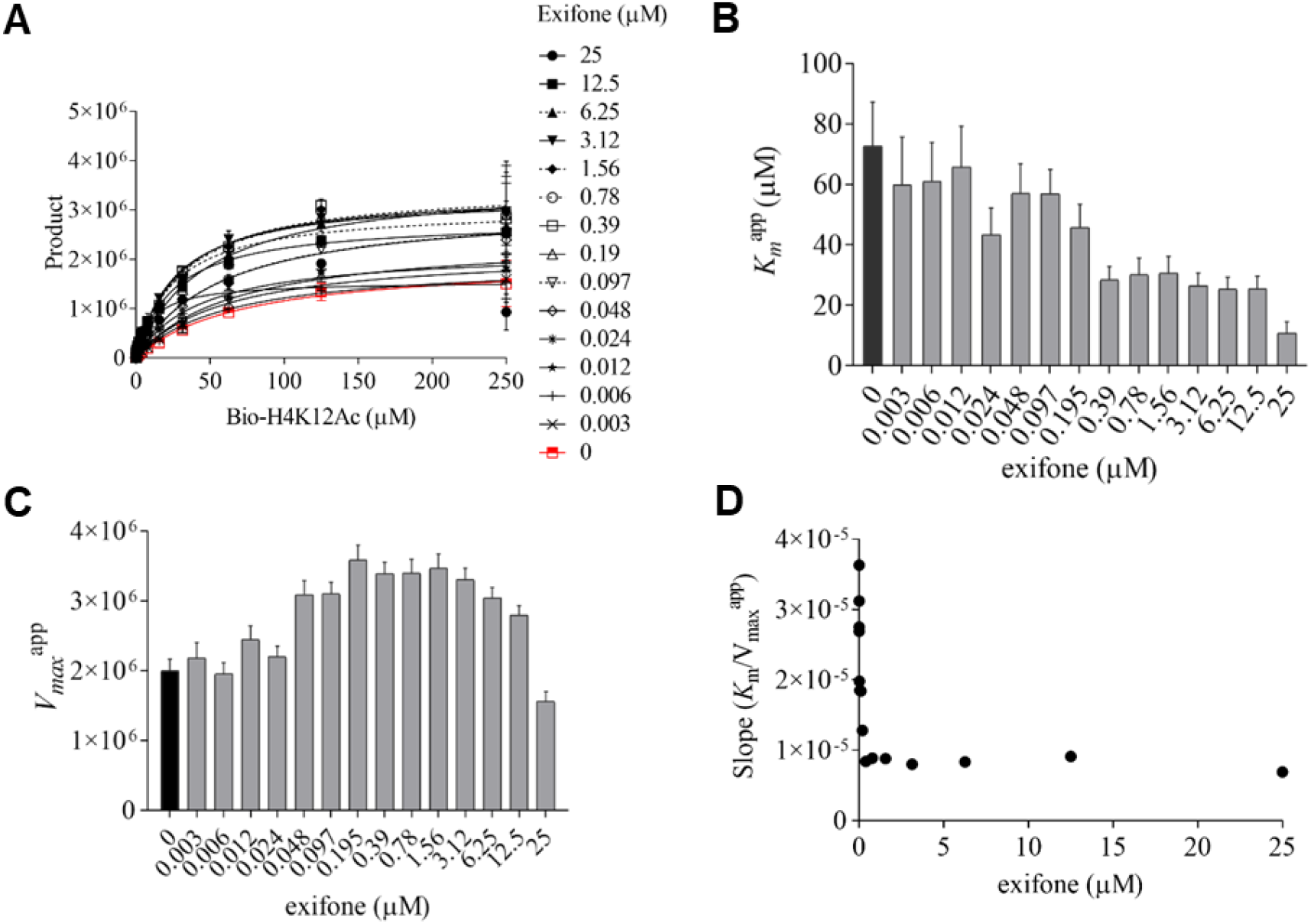
Exifone’s mechanism of action with HDAC1 is consistent with mixed non-essential activation. (**A**) Reaction mechanism of HDAC1 activation by exifone as determined by varying concentrations of both substrate and activator. Increasing exifone concentration decreased the apparent substrate *K*_m_ for Bio-H4K12Ac peptide **(B)** and increased the *V*_max_^app^ (**C**). Replot of slope (ratio of *K*_m_/*V*_max_) vs. exifone concentration shown as descending hyperbolic curve (**D**).

We observed that exifone increased the rate of histone deacetylase reaction. That is, the apparent *V*_max_ was seen to increase with increasing concentration of the HDAC1 activator exifone (**Figure 2C**). With further analysis, the slope was estimated as a ratio of *K*_m_ and apparent *V*_max_ at different exifone (activator) concentrations (**Figure 2D**). Replots of slope vs. activator concentration resulted in a descending hyperbola that is the characteristic of non-essential activation, in which the reaction can occur in the absence of the activator. Non-essential activation can be analyzed in the same manner as partial or mixed-type inhibition, but the changes are in the opposite direction^21^. Thus, exifone acts as a mixed, non-essential activator of HDAC1 and is capable of binding to both free and substrate-bound enzyme. Decreasing substrate *K*_m_ value and increasing *V*_max_ value as a function of activator concentration demonstrates exifone’s influence in Bio-H4K12Ac substrate binding and catalysis.

### Exifone is capable of partially reversing the inhibitory effect of CI-994 in a concentration-dependent manner

Dose-dependent inhibition of HDAC1 by the active site inhibitor CI-994 was investigated in the presence and absence of the activator exifone. CI-994 alone shows an IC50 value of 0.037 μM for HDAC1, and the top fitted value is less than 100% under these test conditions (**Figure 3A**). When HDAC1 was pre-incubated for 15 minutes with 1 μM or 10 μM exifone, the top values for deacetylase activity reached 620% and 450%, respectively, in the dose-response curve for CI-994. These results indicate that the pre-incubation of HDAC1 with exifone can still lead to an increase of HDAC1 catalytic activity at the lower concentrations of CI-994 that were tested.

**Figure 3.**
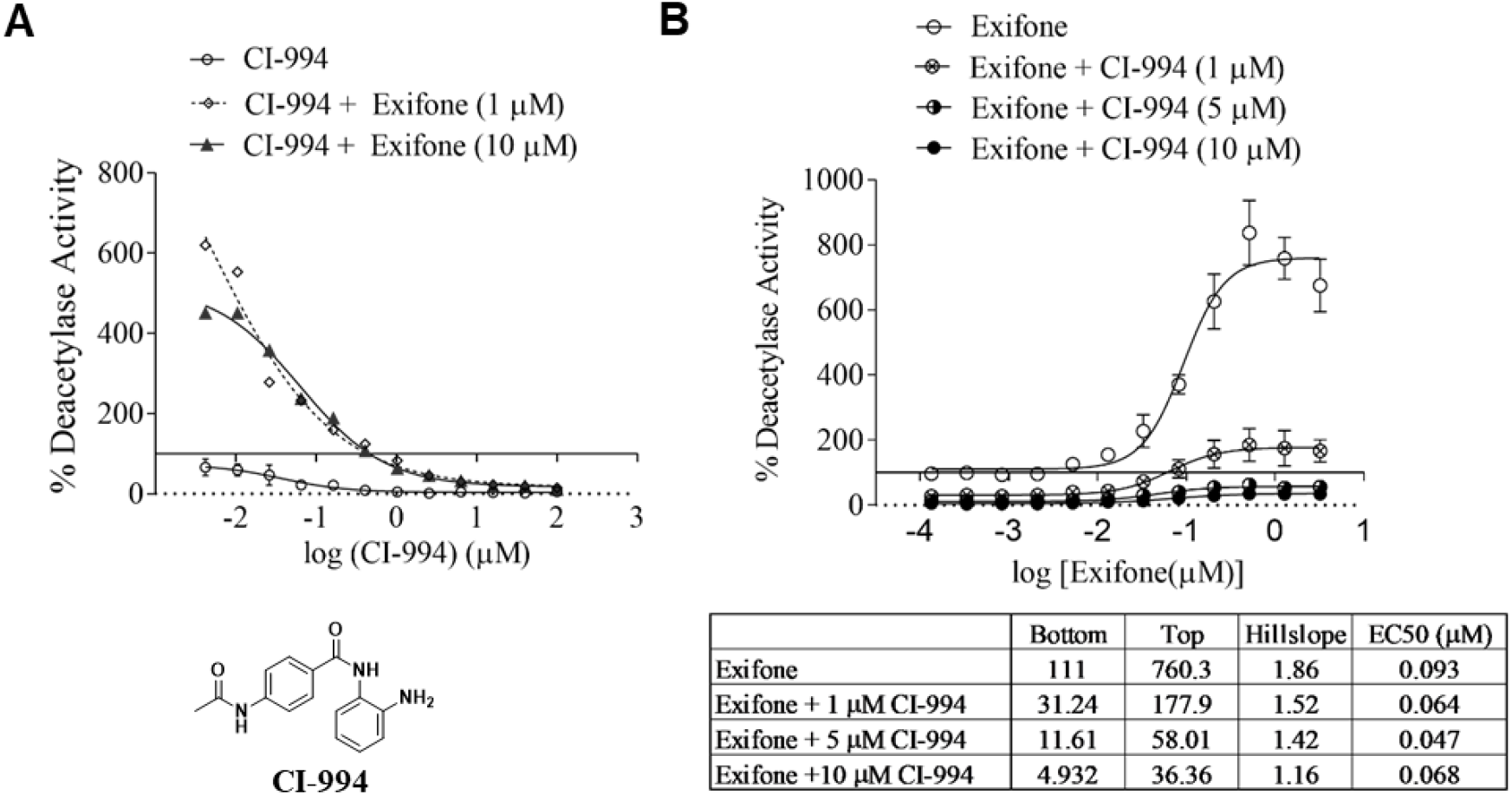
Exifone is capable of reducing inhibition caused by pre-incubation of HDAC1 with active site inhibitor CI-994. (**A**) Concentration response curves for the HDAC inhibitor CI-994 in the absence and presence of the HDAC activator exifone. CI-994 shows an IC50 value of 0.037 μM and the top fitted value is less than 100%. Preincubation of HDAC1 for 15 minutes with 1 μM or 10 μM exifone resulted in the top values for % deacetylase activity reaching up to 620% and 450% respectively. The results indicate that preincubation of exifone can still result in significant HDAC1 activation at lower concentrations of CI-994. (**B**) A dose-dependent increase in HDAC1 activity was observed even if HDAC1 was pre-incubated for two hours with 1 μM CI-994, 5 μM CI-994, 10 μM CI-994, as shown by the best-fit values.

To investigate further, HDAC1 was pre-incubated for two hours with 1 μM, 5 μM, and 10 μM of the active site inhibitor CI-994, followed by an assay to determine the dose-effect of the HDAC1 activator exifone. Exifone shows an EC_50_ value of 0.045 μM, with maximal activation values up to nine-fold when HDAC1 was pre-incubated with exifone at room temperature for two hours before adding the substrate peptide to the reaction mix. When HDAC1 was pre-incubated with 1 μM CI-994, the dose dependence curve for exifone was fitted to a deacetylase activity bottom value of 31.24 and a top value of 177.9. Therefore, exifone was capable of reversing inhibition caused by the pre-incubation of HDAC1 with CI-994 at 1 μM concentration. However, when the enzyme was pre-incubated with 5 μM or 10 μM CI-994, inhibition was no longer reversed by exifone (**Figure 3B**). Pre-incubation studies with the HDAC active-site inhibitor CI-994 at low concentrations demonstrate that exifone is capable of partially reversing inhibition caused by CI-994, indicating interaction directly or indirectly with residues of the HDAC1 active site.

### Exifone displays a preference for activating HDAC1 over HDAC2 in vitro

The concentration response curve for exifone was determined with HDAC1 and its structurally related isoform HDAC2 using Bio-H4K12Ac (1 μM) as the substrate. Top fitted values of HDAC1 vs. HDAC2 activity in dose-response curves to exifone, expressed as a percentage of HDAC activity, were 495.6 and 410.7, respectively. The EC_50_ value for exifone was 0.02 μM for HDAC1, in contrast to 0.08 μM for HDAC2. Estimating the top fitted values as an approximation of relative maximal velocity (*RV*_max_), EC1.5 was calculated as described by Dai et al. (2010)^22^. EC1.5 values of exifone for HDAC1 and HDAC2 were 0.002 μM and 0.015 μM, respectively (**Figure 4**). At least five times more exifone would be required for achieving a similar level of HDAC2 activation, relative to HDAC1. Therefore, exifone appears to be more efficient and show preferential activation of HDAC1. HDAC1 selectivity results revealed that exifone is at least four-fold selective for activating HDAC1, indicating a positive influence on the histone deacetylase isoform that is known to associate with neuroprotection.

**Figure 4.**
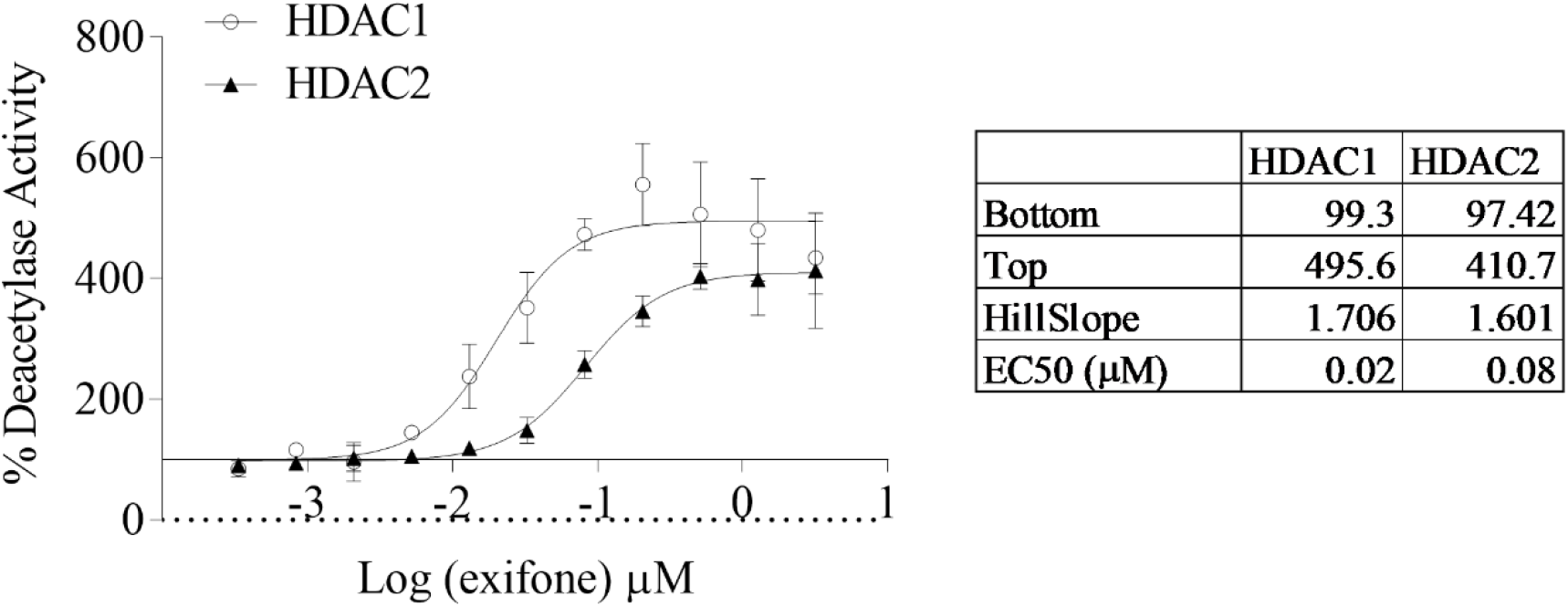
Exifone is at least four-fold selective for activating HDAC1 when compared with HDAC2. The data were fitted with the equation for sigmoidal doseresponse (variable slope) to estimate the EC_50_ values for exifone, and the best fit values are shown as calculated using the RapidFire mass spectrometry assay. The observed EC_50_ values under the defined conditions for HDAC1 are 0.02 μM as compared with 0.082 μM for HDAC2. Estimated EC_1.5_ values for HDAC1 and HDAC2 for this set of data were 0.002 μM and 0.015 μM respectively.

### Biophysical evidence and binding kinetics confirm HDAC1 as a preferred target of exifone

As a small, polyphenolic molecule, exifone is expected to interact with multiple biological targets^23^, such as structurally related class I HDACs. We aimed, therefore, to investigate how exifone interacts with HDAC1 using biophysical assays to both test whether a direct binding interaction can be detected as an assay orthogonal to the mass spectrometry enzymatic assays and to establish a means of determining whether exifone shows preferential targeting of HDAC1 relative to other potential targets relevant to neurodegeneration based on binding kinetics.

Binding kinetics of exifone was first determined using biolayer interferometry (BLI) with biotinylated recombinant HDAC1, HDAC2, and HDAC8. We observed that exifone could bind HDAC1 (**Figure 5A, 5B**), HDAC2 (**Supplementary Figure 2A, 2B**), and HDAC8 (**Supplementary Figure 3A, 3B**). Following reference subtraction, the processed graphs show a distinct pattern of association and dissociation in response to increasing concentrations of exifone. More significantly, benzophenone, failed to show a binding response to HDAC1 in the BLI assay (**Figure 5C**), consistent with its lack of HDAC1 enzymatic activation. Additionally, the BLI results on the interaction of exifone with HDAC1 and HDAC2, indicate a modest preference for HDAC1 binding based on the higher association rate constant (*K*_a_) value and a 1.9-fold lower value for the equilibrium dissociation constant (*K*_D_) with the estimated *K*_D_ values of 0.093 μM and 0.142 μM for HDAC1 and HDAC2, respectively.

**Figure 5.**
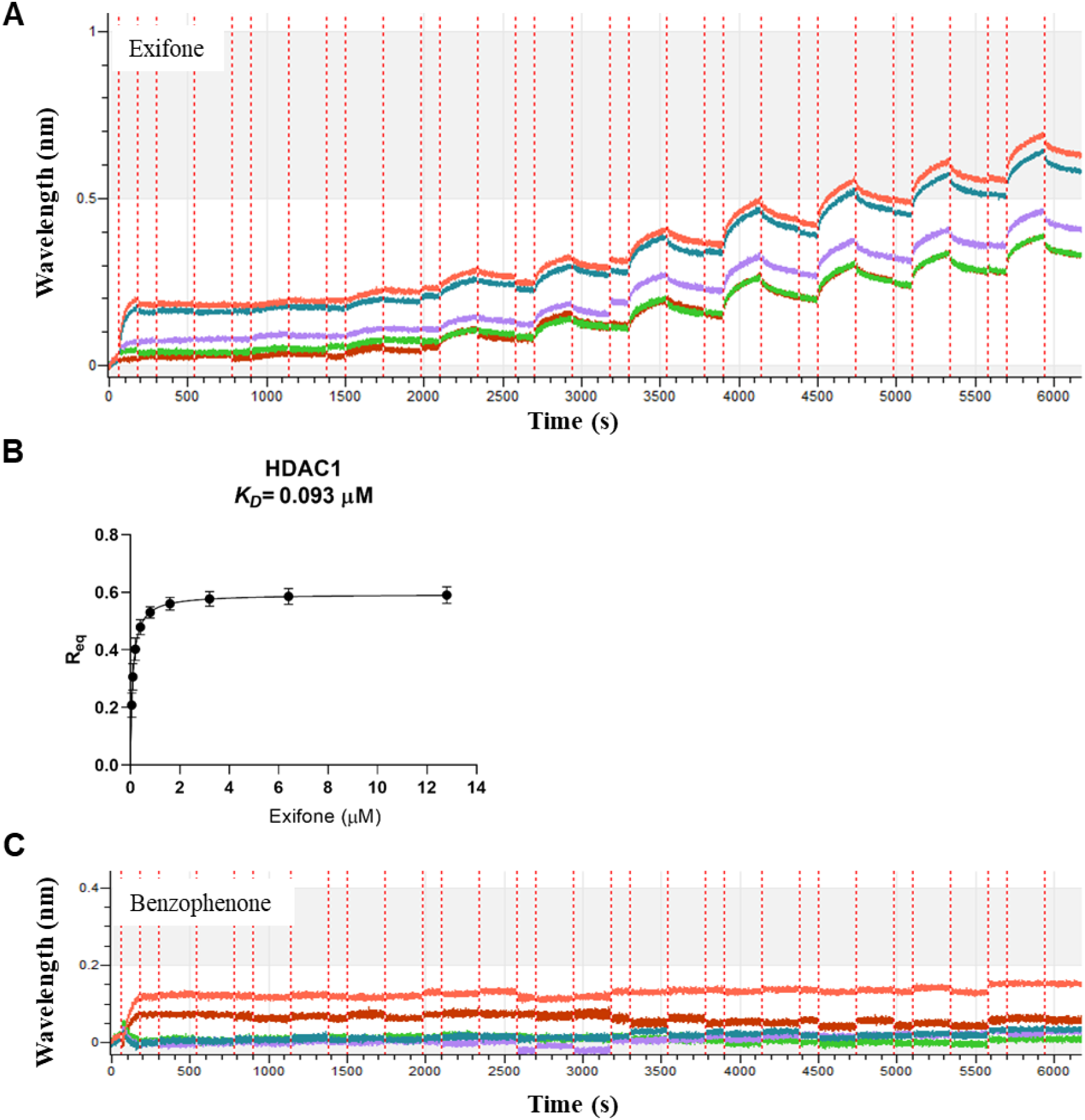
Biolayer interferometry-based biophysical assays of exifone binding to HDAC1. (**A** and **B**) Exifone directly binds HDAC1 as measured using BLI whereas the structurally simpler analog, benzophenone does not exhibit a binding interaction. (**C**). The processed data graphs were obtained after reference (n=3) subtraction and show the acquisition of real-time data in a binding kinetics experiment. After an initial baseline step in the assay buffer, streptavidin biosensors were dipped into solution with the biotinylated HDAC1 for the loading step. Subsequently, a second baseline was performed, followed by association and dissociation of the small molecule analyte (n=5) in solution. Successful binding interaction results in distinct spectral shift pattern that is represented on the sensorgram as a change in the wavelength (nm shift). The data were fit globally with the 1:1 binding model and a representative plot of R_eq_ vs. exifone concentration for the estimation of *K*_D_ is shown (**B**).

Since polyphenolic small molecules like exifone are known to interact with multiple kinase targets, we next sought to determine if exifone also interacted with CDK5^23^, a key regulatory enzyme that when hyperactivated due to binding to the p25 regulatory subunit that is generated by calpain-mediated cleavage of p35 leads to HDAC1 inhibition and elevated DNA damage and neurodegeneration. Using a BLI assay format again, exifone was found to bind to CDK5/p25 with an estimated *K*_D_ value of 0.24 μM (**Figure 6A**), which means based on the measured binding affinities, exifone shows at least a 2.4-fold preference for HDAC1 (**Figure 6A**).

**Figure 6.**
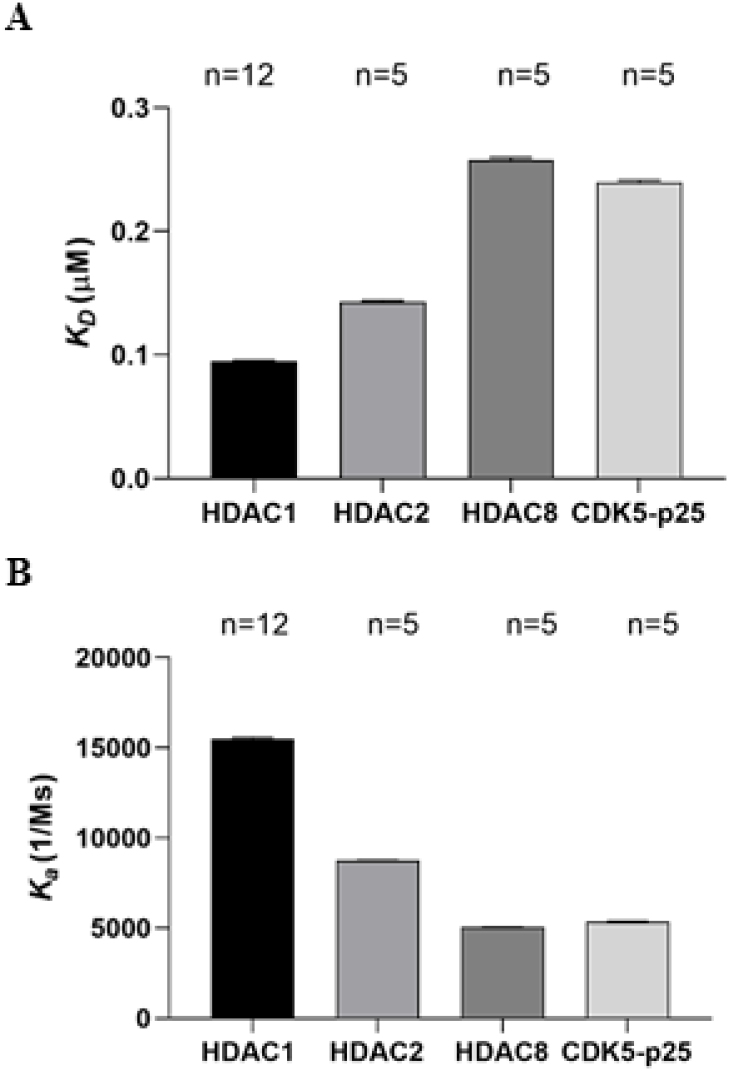
Selectivity profiling of exifone against neurodegeneration relevant targets. Kinetic parameters estimated via BLI assays indicate exifone shows a preference interaction with HDAC1. Exifone has the highest affinity for HDAC1 when compared with HDAC2, HDAC8, and CDK5/p25 as evidenced by (**A**) the lowest value of equilibrium dissociation constant, and (**B**) the highest value for the association rate constant (*k*_a_) that represents the number of complexes formed per second between the small molecule and the biotinylated protein in a 1 Molar solution.

To further investigate the interaction of exifone with potential biological targets, a series of additional kinetic parameters were estimated by a global fit of the BLI data (**Table 1**). Notably, exifone showed the highest value for the association rate constant (*k*_a_) with HDAC1 when compared with the other targets tested (**Figure 6B**), where the *k*_a_, in this case, represents the number of complexes formed per second between exifone and the biotinylated protein in a 1 M solution. Altogether, these observations are significant in establishing that HDAC1 as a preferential target amongst the set of proteins analyzed, although interaction with other targets cannot be excluded from these data alone.

**Table 1.** Summary of kinetic parameters for exifone determined by a global fit of the BLI data. Values calculated from either n=12 replicates (HDAC1) or n=5 replicates (HDAC2, HDAC8, CDK5/p25).

| Parameter | HDAC1 | HDAC2 | HDAC8 | CDK5/p25 |
| --- | --- | --- | --- | --- |
| **$K_D$ (μM)** | **0.0949** | **0.1428** | **0.2579** | **0.2398** |
| $K_D$ Error | 0.0008 | 0.0010 | 0.0017 | 0.0017 |
| **$k_a$ (1/Ms)** | **1.55E+04** | **8.74E+03** | **5.05E+03** | **5.36E+03** |
| $k_a$ Error | 6.49E+01 | 2.55E+01 | 1.39E+01 | 1.52E+01 |
| **$k_{dis}$ (1/s)** | **1.34E-03** | **1.25E-03** | **1.30E-03** | **1.29E-03** |
| $k_{dis}$ Error | 1.11E-05 | 8.09E-06 | 7.90E-06 | 8.10E-06 |
| **$k_{obs}$ (1/s)** | **1.94E-03** | **1.69E-03** | **1.56E-03** | **1.55E-03** |
| $k_{obs}$ Error | 1.34E-05 | 9.36E-06 | 8.59E-06 | 8.86E-06 |
| **Full $R^2$** | **0.8380** | **0.8396** | **0.8817** | **0.8645** |

### Exifone acts as a Deacetylation Activator in Human Neural Progenitor Cells

Following our demonstration of exifone as a potent small molecule that can increase the catalytic activity of HDAC1 *in vitro* in biochemical assays and high-affinity interactor in biophysical assays, we proceeded next to evaluate the deacetylase activating ability of exifone in an *ex vivo* neuronal context using human iPSC-derived neural progenitor cells (NPCs). Previously, we reported the use of mouse primary cortical neurons or human iPSC-derived neurons to examine the cellular activities of HDAC inhibitors^24–26^. We established that the inhibition of HDAC enzymes leads to the increase of the acetylation level of the histone mark H3K9 (H3K9Ac). We expected the activation of HDAC1 by exifone to decrease the level of H3K9Ac. To test this hypothesis, we treated NPCs with exifone (0.5 μM and 2 μM) for 6h or 18h followed by immunostaining of H3K9Ac and quantification of signal from nuclei. Treatment with exifone led to a decrease of the H3K9Ac levels in NPCs (**Figure 7**), providing evidence that exifone can act as an activator of histone deacetylation in an *ex vivo* neuronal setting.

**Figure 7.**
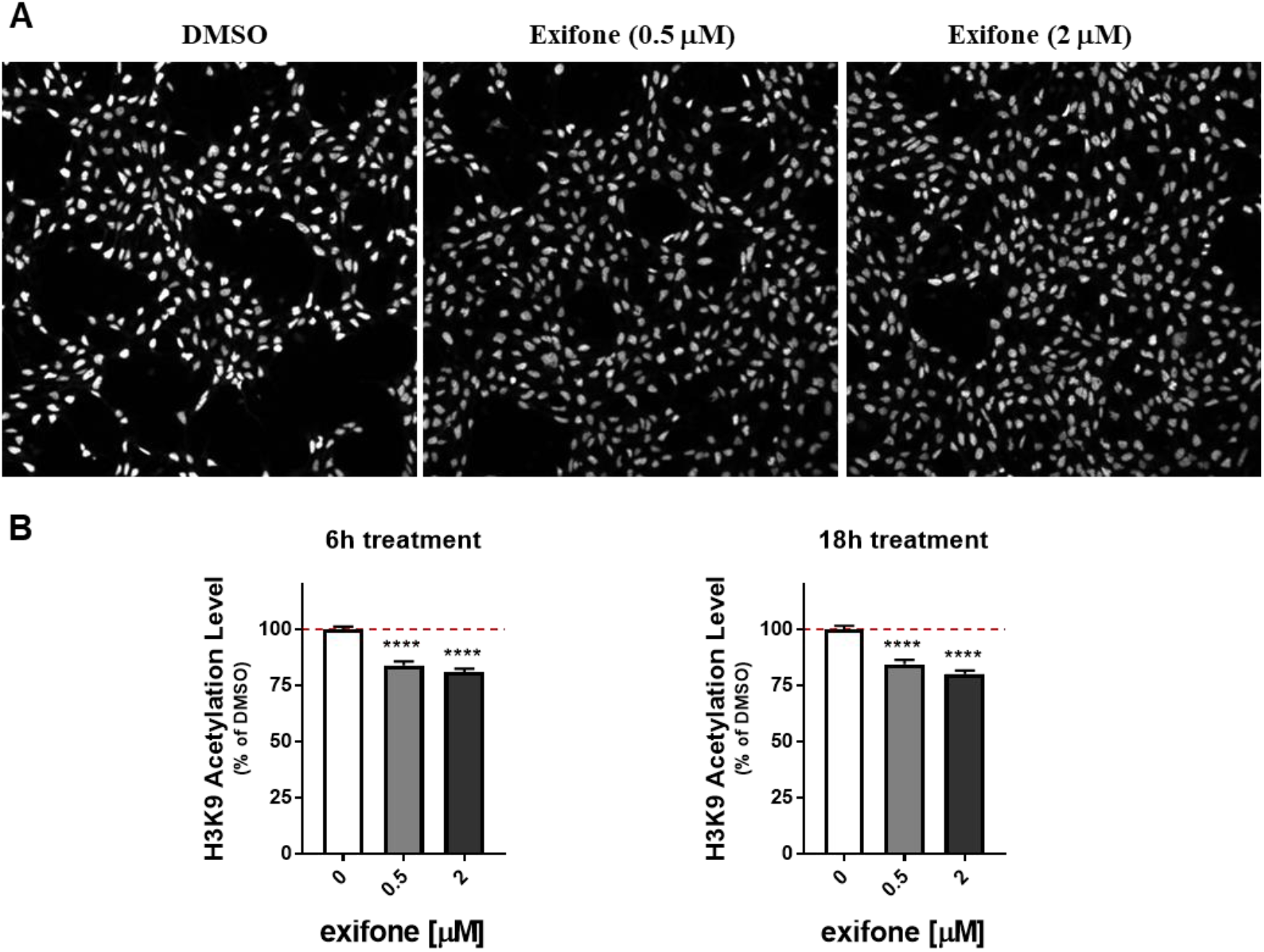
Exifone as an HDAC1 activator increases histone deacetylation in human neural progenitor cells (NPCs). NPCs were treated with various concentrations of exifone for 6h or 18h, followed by fixation and immunostaining for H3K9Ac in nuclei. (**A**) Representative images of H3K9Ac immunostaining of iPSC-derived NPCs, showing dimmer nuclei in 0.5 μM and 2 μM exifone-treated (6h) cultures. Quantitative image analysis was performed to determine the intensity of H3K9Ac in the nuclei. (**B**) Quantitation of the H3K9Ac intensity levels indicates effective activation of HDAC(s) in NPCs, thereby leading to decreased H3K9Ac levels in 6h or 18h treated cultures. Error bars represent standard error of the mean (SEM) based on N=3 biological replicates per treatment. Mean pixel intensities were collected from 9 images per biological replicate totaling approximately 3,500 nuclei. Unpaired t test: **** p < 0.0001.

Our *in vitro* characterization of exifone as an HDAC activator indicates that exifone can also have an effect on other HDACs, including HDAC2, although weaker than on HDAC1 (**Figure 4** and **Figure 6**). This phenomenon was potentially reflected in the cellular acetylation assay, in which the H3K9Ac levels were decreased most effectively by exifone at 0.5 μM and 2 μM, whereas the decrease was attenuated by exifone treatment at 5 μM or 10 μM (**Supplementary Figure 7**). The amount of decrease of H3K9Ac levels became smaller with exifone at 5 or 10 μM, suggesting a loss of specificity on HDAC1 activation by exifone at the higher concentrations.

### Exifone Rescues Neuronal Viability in FTD-Tau Neurons

Patient iPSC-derived neuronal cell models can recapitulate specific disease-relevant phenotypes and are increasingly recognized as robust systems for drug discovery^27^. To evaluate the neuroprotective capabilities of exifone in a neurodegenerative disease-relevance context, we employed an iPSC-derived neuronal cell model from an FTD patient harboring a tau-A152T variant that shows an early accumulation of tau protein in the form of phospho-tau species of reduced solubility, and as a consequence exhibits selective vulnerability to stress^28^. This phenotype observed in patient-derived neurons is characterized by >60% neuronal death upon treatment with the stressor rotenone that inhibits the mitochondrial electron transport chain (ETC) complex I (**Figure 8**). In contrast, rotenone leads to <10% neuronal death in control non-mutant neurons (**Figure 8**).

**Figure 8.**
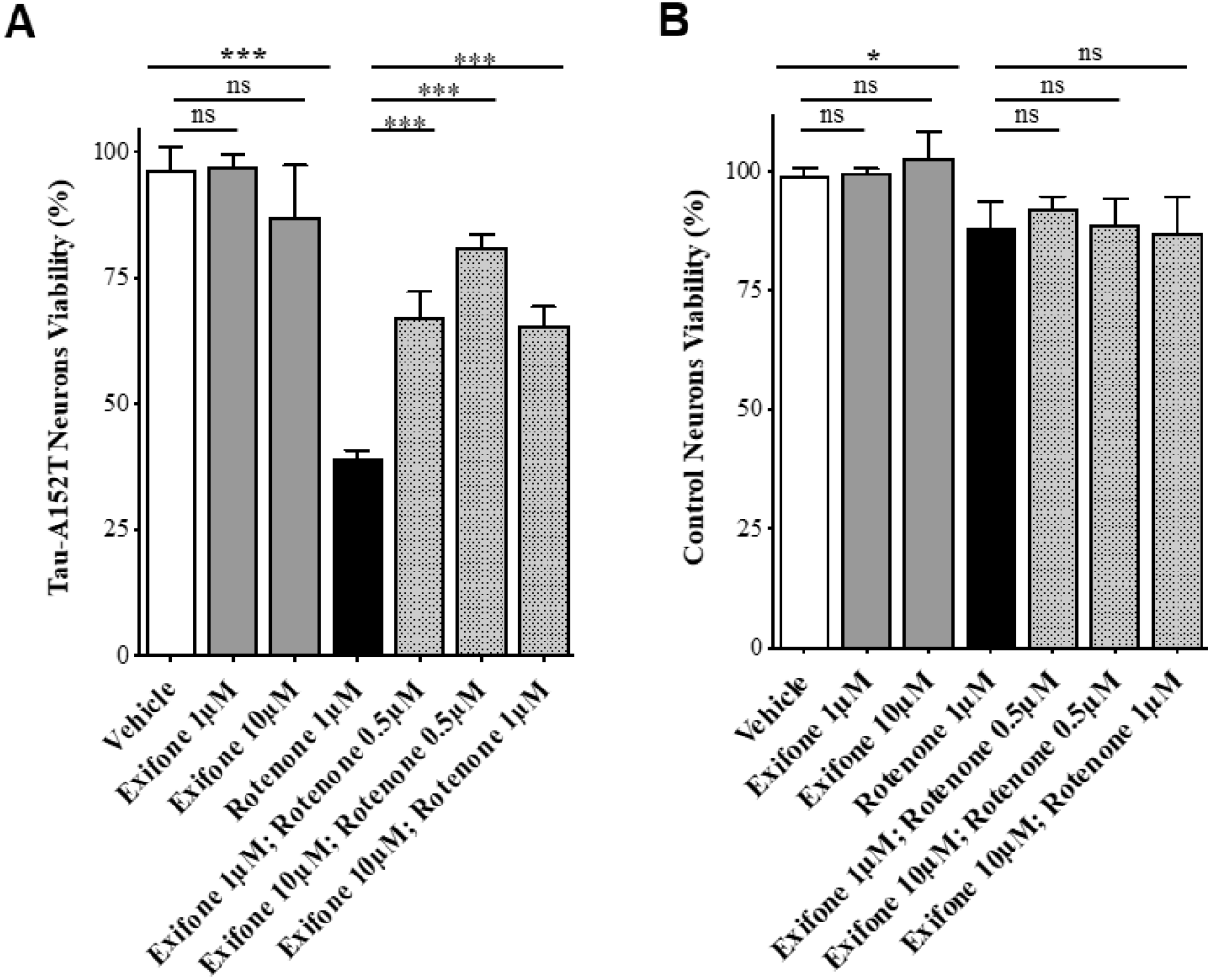
Exifone rescues stress vulnerability of FTD tau-A152T neurons. Cell viability of tau-A152T (**A**) or control (**B**) neurons (differentiated for eight weeks from NPCs), treated with compounds for 24h ±pre-treatment with exifone (8h). Values represent % viability (±SD) relative to vehicle-treated neurons. Student t-test \**p*<0.05, \*\*\**p*<0.001, ^ns^*p*> 0.05 (*n*=2 with technical triplicates).

To test whether exifone can protect neurons against tau-mediated stress vulnerability, we pre-treated neurons with 1 μM or 10 μM of exifone for 8h, followed by the addition of stressor rotenone at 0.5 μM or 1 μM of dose (for a total of 24h treatment). Exifone alone did not affect viability of the control or FTD neurons (**Figure 8**), but when added to the neuronal cultures before stress induction by rotenone, exifone rescued cell viability to almost 80% in a dose-dependent manner (**Figure 8**). To determine if the effect of exifone in tau-A152T neuronal viability was the result of direct modulation of tau levels, we examined levels of total tau and phospho-tau (P-tau-S396) by western blot upon 24h treatment with 10 μM exifone. No significant downregulation of tau by exifone was detected (*data not shown*) relative to vehicle-treated cells. These results suggest that exifone may have a protective effect in human neurons against neurodegeneration.

## Discussion

Neuroprotection represents a broad set of molecular and cellular mechanisms that can promote rescue, recovery, and regeneration of the structure and function of neuronal cells^29^. Defining targets regulating these mechanisms if of great interest from the perspective of gaining fundamental insight into nervous system function and potential therapeutic applications. In these contexts, activation of class I HDACs via small molecules has been the subject of only a limited number of studies^4, 30-31^. With the initial discovery of the role of an increase of HDAC1 activity in the reduction of neurotoxicity due to aberrant activity of CDK5/p25^1^, our investigations have resulted in the identification of multiple small-molecule HDAC1 activators^4, 32^ In particular, by pursuing the structure-activity relationship of a series of synthetic HDAC1 activators, exifone was identified as a potent activator of HDAC1 with nanomolar potency in RapidFire MS assays using acetylated substrates derived from both histone (H4K12Ac) and non-histone (p53K382Ac) sequences^32^. Here our use of a mass spectrometry-based assay (RapidFire MS) and orthogonal biophysical assays based upon BLI, rather than relying on assays with non-physiological, fluorophore-modified peptides, was designed to overcome challenges encountered previously when developing activators of a structurally distinct family of NAD^+^-dependent lysine deacetylases^19–20^.

In the present study, the mechanism of HDAC1 activation by exifone appeared to be consistent with non-essential activation that is usually analyzed in a manner similar to mixed inhibition with the changes in the enzyme activity in the opposite direction^21^. To understand the mechanism further, we carried out inhibition studies with CI-994 [4- acetamido-*N*-(2-aminophenyl)benzamide)], which is a relatively selective inhibitor of HDAC1 and HDAC3^33–34^. Pre-incubation experiments with a low concentration (1 μM) of the CI-994, demonstrated that exifone was capable of partially reversing inhibition caused by CI-994. Inhibition studies with this active site inhibitor CI-994 showed that exifone is likely to interact with sites other than the active site, consistent with evidence for a mixed mechanism of activation. Our results also demonstrate that exifone may have a protective effect in patient-derived human neurons against neurodegenerative stress. As a potent activator of HDAC1, exifone provides a new chemical tool to dissect the function of HDAC1 in epigenetic regulation and regulation of the acetylome. However, there are still several open questions as to the mechanism of HDAC1 activation that will require further investigation to validate these findings, including the elucidation through structural studies of a defined binding site of exifone and related activators and determining how the catalytic properties of HDAC1 are affected. Furthermore, by analogy to the NAD^+^-dependent deacetylase activator studies^19–20^, beyond the histone (AcH4K12-based) and non-histone substrates (Acp53K382-based) tested here, activators of the Zn^2+^-dependent HDAC1 may also turn out to have specificity for certain substrates over others. Thus, working backwards from compounds like exifone that show neuroprotective effects *in vivo* it may be possible then to discern the relevant substrates and develop next-generation HDAC1 activators specific for certain substrates as novel approach to therapeutic development.

Despite having a close structural similarity, HDAC1 and HDAC2 are characterized by distinct functions in the brain^35^. In biochemical assay, we have demonstrated that exifone is at least four-fold selective for activating HDAC1 when compared that with HDAC2; thus, exifone is likely to influence HDAC1 preferentially. Additional evidence from biophysical assay confirmed a modest preference (1.9-fold) for the *in vitro* binding with HDAC1 based on the lower equilibrium dissociation constant value (1.9-fold) compared to HDAC2. More significantly, here we show that exifone can act as an activator of deacetylation in human neural progenitor cells. HDAC1 has been shown to play a vital role in the regulation of neuronal viability^1, 36^. In the present study with FTD patient-derived neurons^28^, exifone treatment alone did not affect viability of the control or FTD neurons (**Figure 8**), but when added to the neuronal cultures before treatment with rotenone, a stressor of the mitochondrial electron transport system, exifone rescued neuronal viability to approximately 80% in a dose-dependent manner (**Figure 8**). This is the first report of a small-molecule HDAC activator exerting biochemical effects in a neuronal setting. Additionally, while the results from our biochemical and biophysical assay panel indicate that HDAC1 is preferentially targeted by exifone over other targets that were evaluated, including HDAC2, HDAC8, and CDK5/p25, given the nature of polyphenols, we cannot rule out the contributions of other targets to the effects of exifone in a cellular or in *vivo* context.

Besides the studies reported here with exifone, HDAC activity has been shown to catalytically activated *in vitro* with a similar mixed mechanism by metabolites of intermediary metabolism^30^. These multiple intracellular metabolites were shown to directly increase the catalytic activity of HDAC1 toward acetylated histones with the EC_50_ values ranging from high micromolar to millimolar range^30^. The molecules that cause *in vitro* activation of recombinant HDAC1 and HDAC2 following a mixed activation kinetics include various coenzyme A (CoA) derivatives and the reduced (NADPH), but not oxidized (NADP^+^), form of nicotinamide adenine dinucleotide phosphate. These findings indicate that the metabolic state of a cell might influence the cellular activity of HDACs^30^. Moreover, these results highlight the opportunity to pharmacologically modulate the activity of HDACs via small molecules utilizing the intrinsic metabolic pathways of the cell. Of note, the intracellular metabolites that have been identified as HDAC1 activators through these studies were unable to enhance HDAC1 activity when a synthetic, fluorophore-labeled, acetylated tripeptide substrate (Fluor de Lys) was used instead of histones^30^. These observations provide evidence that HDAC1 activators may affect enzyme-substrate interactions independent of the acetylated lysine and again point to the importance of using appropriate substrates and assays when characterizing small-molecule activators.

Additionally, the regulation of the enzymatic activities of HDACs 1, 2, and 3 that are present in association with other proteins in multiple complexes, by inositol phosphates was reported by Watson et al. (2016)^37^. This study demonstrated an allosteric mode of communication between the inositol binding site and the active site of the enzyme with various inositol phosphates shown to activate HDAC3 with binding affinities ranging from low to high micromolar values. The catalytic activity of HDAC1 that exists in a complex with MTA1 (metastasis-associated protein) was also shown to be positively influenced by various inositol phosphates^37^. The level of activation observed with these natural intracellular metabolites *in vitro* is modestly higher than the basal level. However, these results provide encouraging insight for HDAC activation by potent small molecules such as exifone that show nanomolar potency in *in vitro* assays.

Activation of another class I HDAC8 has also been the subject of few investigations with the identification of N-acetylthiourea derivatives as highly potent and isozyme selective activators of HDAC8^31^. More significantly, HDAC8 activators may serve as possible leads in the therapeutic management of Cornelia de Lange Syndrome (CdLS) spectrum disorder^38–39^. Exifone can also activate HDAC8 (**Supplementary Figure 1**), albeit with a less potent EC_50_ of 0.27 μM, and an EC1.5 value of 0.08 μM in a RapidFire MS assay in comparison to HDAC1. Thus, although exifone appears to be at least 12-fold more selective for HDAC1 relative to HDAC8, exifone could still offer another route for the development of HDAC8 activators.

Despite the fact that polyphenols generally present challenges from a medicinal chemistry point of view and have the potential to interact with diverse targets, the initial preclinical and clinical studies on exifone that led to its registration did not reveal overt toxicity. It was not until months of administration to elderly patients at doses of 200-600 mg/day that reversible hepatoxicity was observed^10–14^. Questions left unanswered include why such a small fraction of patients (reported to be ~1/15,000) exhibited this hepatotoxicity and whether genetic or possibly other drug interactions contributed to this outcome. Furthermore, without a defined target and functional biomarkers to determine optimal exposure levels, it is unclear whether lower doses or alternating dose schedules would provide a means to reduce the risk hepatotoxicity while retaining the clinical benefit that was observed. Finally, given the known heterogeneity of dementia patients, along with knowledge that when exifone was tested in the 1980s this predated the use of either genetic testing or functional biomarkers such as in the case of the Alzheimer’s-type dementia of levels of amyloid by positron emission tomography (PET) imaging or analysis of cerebrospinal fluid Aβ_1-42_ or phospho-tau levels, to stratify patients and select patients for treatment that were at the earliest rather than late stages of the diseases. Thus, by “*testing the right target and right drug at the right stage*”^40^, as is now considered important in a precision medicine framework for neurotherapeutics^41–43^, it is possible that a specific subpopulation of dementia patients would have shown even more improved outcomes and that the dose and schedule of exifone treatment could have been tuned appropriately to maximize benefit and minimize undesired effects outside of the CNS.

As a step toward to addressing these questions with exifone using preclinical mouse models of AD and age-related accumulation of DNA damage, in a companion study we have recently shown that HDAC1 modulates oxidative DNA repair in the aging brain via 8-oxoguanine DNA glycosylase (OGG1), a DNA glycosylase of the base excision repair (BER) pathway that primarily acts on 8-oxoguanine (8-oxoG)^2^. 8-oxoG is regarded as a biomarker of oxidative stress and considered pre-mutagenic due to the potential ability to pair with adenine instead of cytosine^44^ during DNA replication. HDAC1 deficincy leads to increased 8-oxoG levels, which is a type of oxidative DNA damage associated with transcriptional repression. Neurons lacking HDAC1 exhibited increased DNA damage, and importatnly pharmacological activation of HDAC1 by exifone protected against the harmful effects of oxidative DNA damage in the brains of aged wild type and 5XFAD mice^2^. Taken together, further investigation of exifone and other small molecule HDAC1 activators has the potential to facilitate new avenues for drug development with novel small molecule HDAC1 activators as lead compounds for CNS disorders like AD and other human diseases associated with genomic instability.

## Methods

### RapidFire Mass spectrometry (MS) assay for the detection of histone deacetylase activity

A high-throughput mass spectrometry assay with the RapidFire^™^ platform (Agilent Technology, Wakefield, MA) was used for detection of deacetylation activity with the acetylated peptide substrates derived from histone Bio-H4K12Ac (Anaspec #64849) and non-histone Bio-p53K382Ac (Anaspec #65046) sequences (Anaspec, Fremont, CA). Recombinant HDAC1 and HDAC2 enzyme preps were from BPS Biosciences (San Diego, CA). For the determination of dose-dependence curves, test compounds were preincubated with HDAC1 for 15 minutes in an assay buffer containing 50 mM Tris, pH 7.4, 100 mM KCl, and 0.01% Brij-35. Deacetylation reactions were initiated following the addition of the acetylated peptide substrate. Histone deacetylase assay was performed in standard 384 well plates in a 50 μl reaction, and the reactions were terminated by adding 5 μl of 10% formic acid. For the mechanism of action experiments, deacetylation reactions were performed by varying the concentrations of both the substrate and the test compound. The EC_50_ values of individual compounds were estimated by performing deacetylation reactions with 40 nM enzyme and 1 μM of the acetylated peptide substrate. To keep the substrate conversion values under 10%, the enzymatic reaction durations for Bio-p53K382Ac and Bio-H4K12Ac were 45 and 60 minutes, respectively.

For the MassSpec based detection, assay plates were transferred to a RapidFire200 integrated autosampler/solid-phase extraction (SPE) system (Agilent Technologies, Wakefield, MA) coupled with an API4000 tripl*e* quadrupole mass spectrometer (Applied Biosystems, Concord, Ontario, Canada). Additional details for the RapidFire Mass Spec analysis are reported elsewhere ^45–46^.

The percentage of substrate conversion was calculated as a function of product and substrate mass spectrum peak area. [%Substrate Conversion= 100*[Product/(Product+Substrate)]. Percentage of enzyme activation =100* [(MIN-test compound)/(MIN-MAX)], where MINimal (0%) enzyme activity was detected in the presence of a histone deacetylase inhibitor (10 μM SAHA) and MAXimal enzyme activity of HDAC1 reaction in the absence of test compound as 100% activity. MAX and MIN values were estimated as an average of 16 wells of a single column of 384-well plate. Data were analyzed with GraphPad Prism 8. EC^1.5^ was estimated as the concentration of the activator molecule required to achieve 1.5-fold activation^22^. The data on the activation of histone deacetylase by small molecules were analyzed using non-essential enzyme activation^47^.

### Determination of binding kinetics via Bio-layer interferometry (BLI)

BLI assay for the detection of binding interaction with the recombinant enzyme was performed in the Octet Red384 instrument (ForteBio, Fremont, California) with 1X PBS with 0.01% Brij-35 was used as the assay buffer. The recombinant HDAC proteins (BPS Biosciences, San Diego, CA) were biotinylated using EZ-Link NHS-PEG4-Biotinylation Kit, and excess biotin reagent was removed using Zeba spin desalting column following the manufacturer’s recommendation (Thermo Fisher Scientific, Waltham, MA). The biotinylated protein samples to be used as ‘load’ in the BLI experiments were purified in 1X PBS. For BLI, streptavidin (SA) sensors were used to detect the biophysical interaction between the small molecule ligand and the biotinylated proteins. Before the BLI assay, the streptavidin sensors were soaked by dipping in 200 μL of assay buffer in a 96-well Greiner Bio-One Black flat bottom plate (#655209) The assay was performed in a reaction volume of 80 μL in Greiner Bio-One 384 well black flat bottom PP plates (#781209, Greiner, Monroe, North Carolina) with an initial baseline step, followed by loading of 250 nM biotinylated protein. The recombinant protein and the small molecule samples were arranged in a 384-well plate as per a plate map compatible with the 8-channel mode kinetic analysis, where the sensors move from low to high concentration of the small molecule ligand. Further steps included a second baseline (120s), association (240s), and dissociation (240s) for the subsequent cycles. All the sensors were loaded with the biotinylated protein, and three sensors with appropriate concentration of DMSO (comparable to small molecule samples) in the assay buffer were used as the reference.

Data were analyzed using Data Acquisition HT 11.0 software following reference subtraction (an average of three sensors with DMSO in assay buffer) using the 1:1 binding model with a global fit for the replicates (n=5). Global fit assumes complete dissociation of the binding partner (signal will return to zero at an infinite time)^48^.

The equilibrium dissociation constant (KD) was also estimated using data at equilibrium from each available small molecule ligand concentration using the steady-state analysis. The instrument manufacturer (Fortebio, article #137) recommended the steady-state option for analyzing interactions that are either low affinity or with very fast on and off rates affinity or with very fast on and off rates. For steady-state analysis, R equilibrium (Req) was fitted according to the 1:1 binding model with the equation Response= (R_max_*Conc.)/(K_D_ + Conc.).

When the “R equilibrium” option is selected, Fortebio’s software calculated affinity constants based on the Req values determined from the resultant curve fits. In the steady-state analysis, Req is plotted against the small molecule sample concentration to infer the R_max_. *K*_D_ is estimated as the small molecule concentration where 50% of R_max_ is achieved. As per the instrument manufacturer, if all the curves have reached equilibrium, these two sets of values correspond to “Response,” and R_eq_ values should match. Additional descriptions about BLI assay are available on the manufacturer (ForteBio) website.

### Immunofluorescence Analysis of Histone Acetylation Using Human iPSC-Derived Neural Progenitor Cells (NPCs)

Human NPCs were seeded in poly-ornithine/laminin-coated 96-well plates (Corning #3904) at 30,000 cells per well, and treated next day with exifone at 0.5, or 2 μM in triplicate for 6h or 18h followed by fixation and immunostaining for the acetyl-histone H3- Lys9 mark (monoclonal Anti-H3K9Ac; Millipore, #07-352). Nine images at 20X magnification were collected and analyzed from each well from a 96-well plate on an automated confocal microscope, IN Cell Analyzer 6000 (GE Healthcare). Immunofluorescent intensities of H3K9Ac mark in the nuclei (total ~ 2500 – 4500 nuclei/treatment) were quantified by high-content image analysis (IN Cell Analyzer Workstation 3.7.2, GE Healthcare). H3K9 acetylation levels were reported after the intensities of the acetyl mark were normalized to DMSO-treated samples. Unpaired t-tests were used to determine treatment significance for all compounds tested. Stars of significance indicate a significant effect for a treatment dose compared to control (* 0.01 ≤ p < 0.05, ** 0.001 ≤ p < 0.01, *** 0.0001 ≤ p < 0.001, **** p < 0.0001).

### Human iPSC-derived neuronal cultures

Induced pluripotent stem cell (iPSC) lines and derived neural progenitor cell (NPC) lines for the control (8330-8-RC1) and FTD tau-A152T (FTD19-L5-RC6) have been described previously.^28^ Briefly, NPCs were cultured on poly-ornithine, and laminin (POL)-coated plates, with DMEM/F12-B27 media supplemented with the growth factors EGF and FGF, and heparin, and passaged with TrypLE (Life Technologies). Neural differentiation was achieved by plating NPCs at an average density of 50,000 cells/cm^2^ on POL plates, with DMEM/F12-B27 media only (no growth factors), with half-media replacement every three days, for a total of eight weeks.

### Compound treatment and viability assay in neuronal cultures

NPCs were differentiated in DMEM/F12-B27 media, for eight weeks. For testing neuronal viability upon exifone or rotenone (Enzo Lifesciences) treatment, compound, or vehicle alone (DMSO) was added directly to the media. After 24h incubation, cell viability was measured with the Alamar Blue Cell viability reagent (Life Technologies), according to the manufacturer’s instructions. Fluorescence was measured with the EnVision Multi-label Plate Reader (Perkin Elmer). For rotenone-induced stress rescue experiments, eight-week differentiated neurons were pre-treated with either 1 μM or 10 μM of exifone for 8h. Then, the stressor compound rotenone was added directly to the same media for a total of 24h incubation, at which point cell viability was measured as described above.^28^

### Western blot analysis

Neuronal cells were washed and collected in ice-cold PBS by scraping, pelleted at 3000*g* for 5 min, and lysed in RIPA buffer (Boston Bio-Products) supplemented with 2% SDS (Sigma), protease inhibitors (Roche Complete Mini tablets), and phosphatase inhibitors (Sigma). Protein concentrations were estimated with the Pierce BCA Protein Assay Kit (Thermo Fisher Scientific), and Western blot analysis was done using the Novex NuPAGE SDS-PAGE Gel System (Invitrogen) and standard immunoblotting techniques^28^. Membranes were exposed to autoradiographic film (LabScientific), then films were scanned using a GS-800 Calibrated Densitometer (Bio-Rad), and band intensities (pixel mean intensity) were quantified using Adobe Photoshop CS5 Histogram function. Tau band intensities were calculated relative to the respective loading-control actin band. Antibodies were as follows: total tau antibody TAU5 (Invitrogen AHB0042), phospho-tau S396 (Invitrogen 44752G), and loading control β-Actin (Sigma A1978).

## Acknowledgments

We wish to thank Dr. Kelly L. Arnett and Harvard’s Center for Macromolecular Interactions for advice regarding Biolayer Interferometry (BLI) and Dr. Peter Rye, Agilent Technologies, for advice regarding RapidFire MS. We also wish to thank Dr. Kenneth S. Kosik, UC Santa Barbara, for providing us the recombinant CDK5/p25 preparation. This work was supported by a NIA grant (RC1 AG035711) to L.-H.T./S/J.H., an Alzheimer’s Association New Investigator Research Program (S.J.H.), the Tau Consortium/Rainwater Foundation (S.J.H). and the Stuart & Suzanne Steele MGH Research Scholar award to S.J.H. L.-H.T. also received funding from NIA Grant (AG046174), NINDS Grant (NS102730), and Glenn Foundation Award for research in biological mechanisms of Aging.

## SUPPLEMENTARY INFORMATION

**Supplementary Table 1.** The bottom and top fitted values from the dose response curves (Figure 1H) represent % deacetylase activity and activation of HDAC1 by exifone at variable concentrations of the acetylated substrate peptide (Bio-H4K12Ac). The highest level of enzyme activation was observed when the substrate concentration is significantly lower than the apparent *K*_m_ value for the acetylated peptide substrate ([S]<<[K_m_]).

| Bio-H4K12Ac ($\mu\text{M}$) | Bottom | Top | EC50 ($\mu\text{M}$) |
| --- | --- | --- | --- |
| 62.50 | 102.10 | 212.2 | 0.063 |
| 31.25 | 106.90 | 257.7 | 0.053 |
| 15.60 | 109.20 | 294.9 | 0.081 |
| 7.80 | 92.29 | 299.2 | 0.077 |
| 3.90 | 94.33 | 352.4 | 0.10 |
| 1.95 | 89.24 | 362.5 | 0.10 |
| 0.97 | 79.05 | 540.3 | 0.16 |
| 0.48 | 46.27 | 561.3 | 0.15 |
| 0.24 | 91.39 | 990.8 | 0.23 |
| 0.12 | 12.33 | 2613 | 0.19 |

**Supplementary Figure 1.**
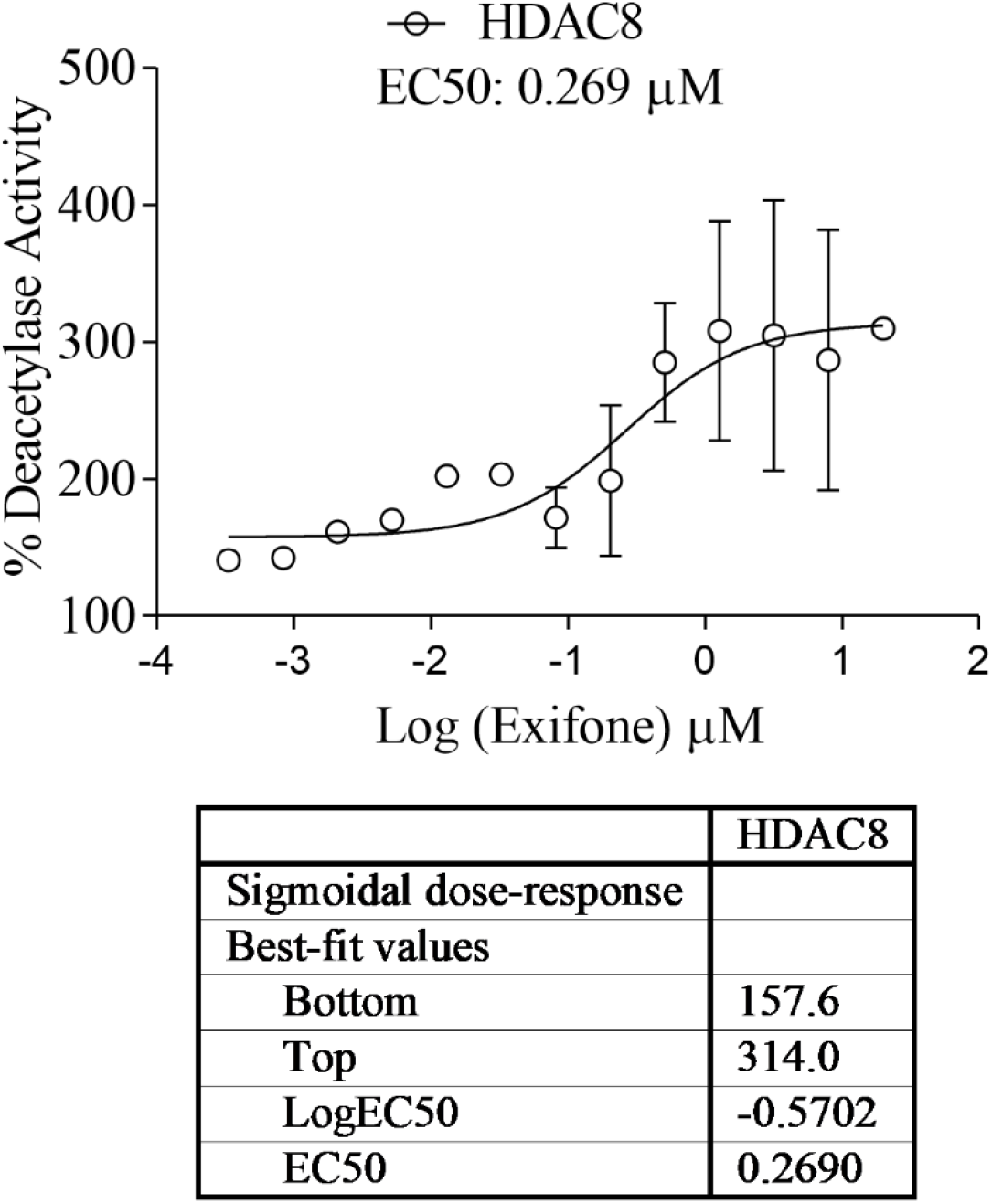
Exifone can increase the rate of deacetylation catalyzed by HDAC8 and shows an EC_50_ value of 0.27 μM and EC_1.5_ value of 0.08 μM.

**Supplementary Figure 2.**
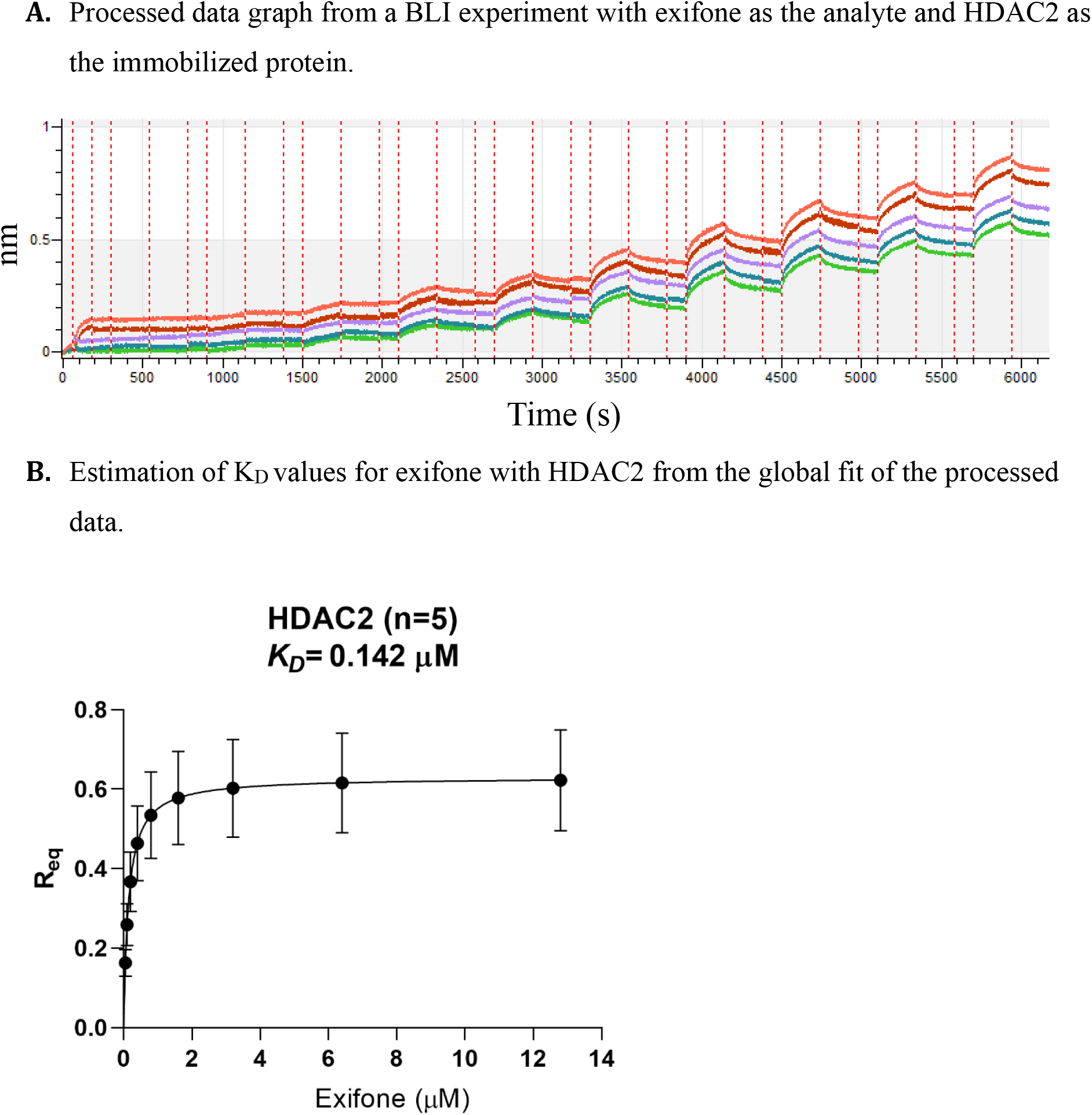

**Supplementary Figure 3.**
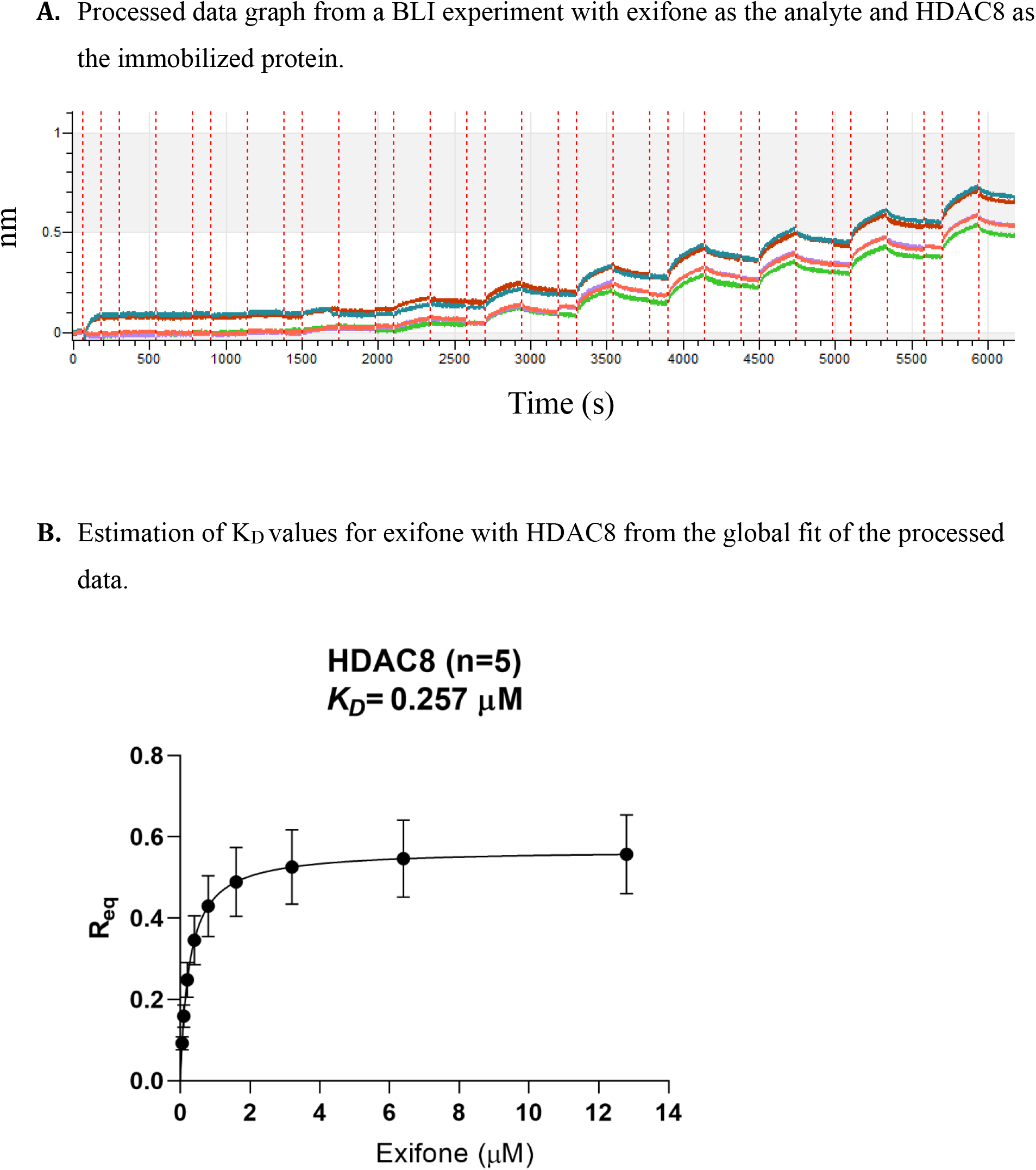

**Supplementary Figure 4.**
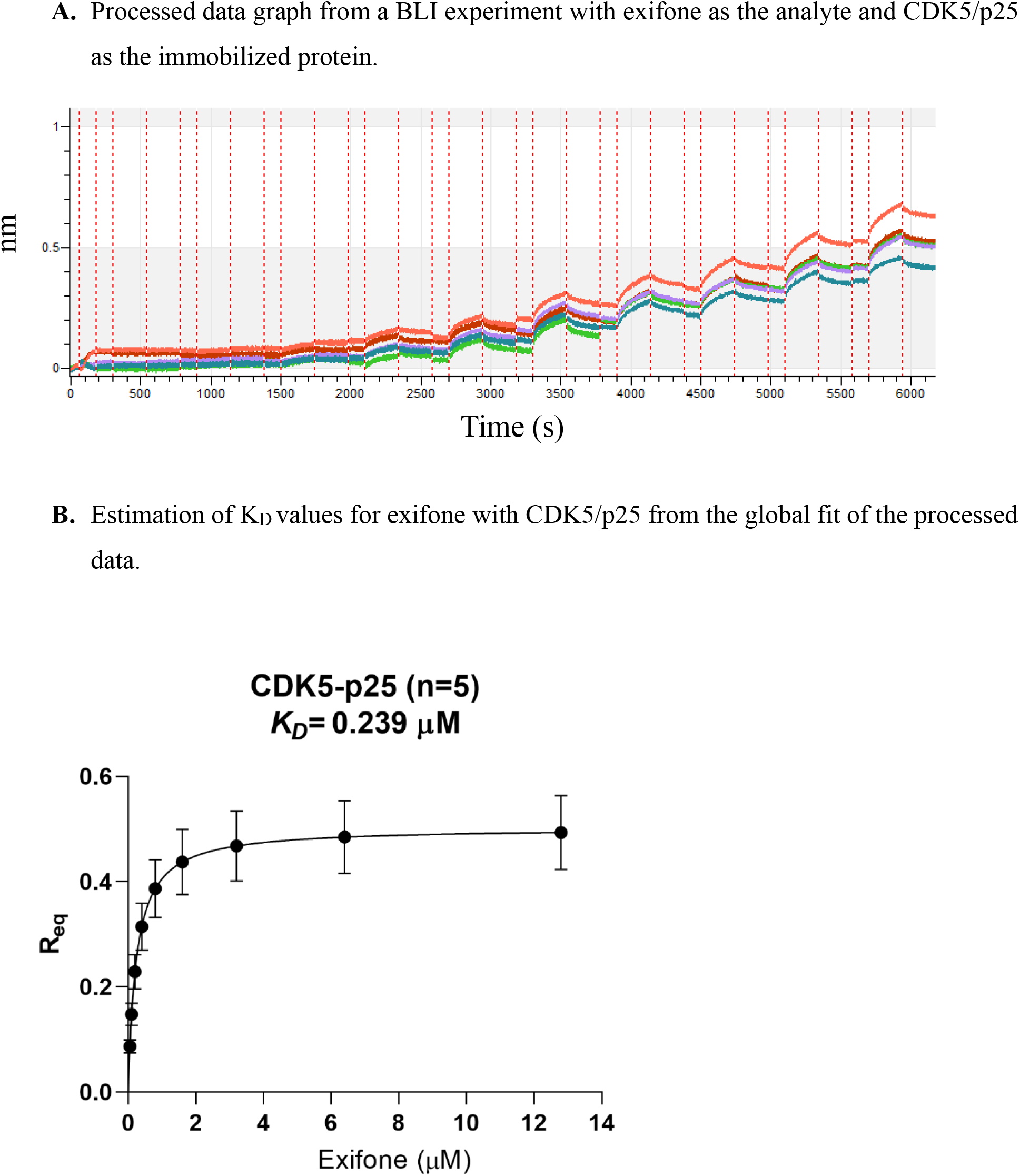

**Supplementary Figure 5.**
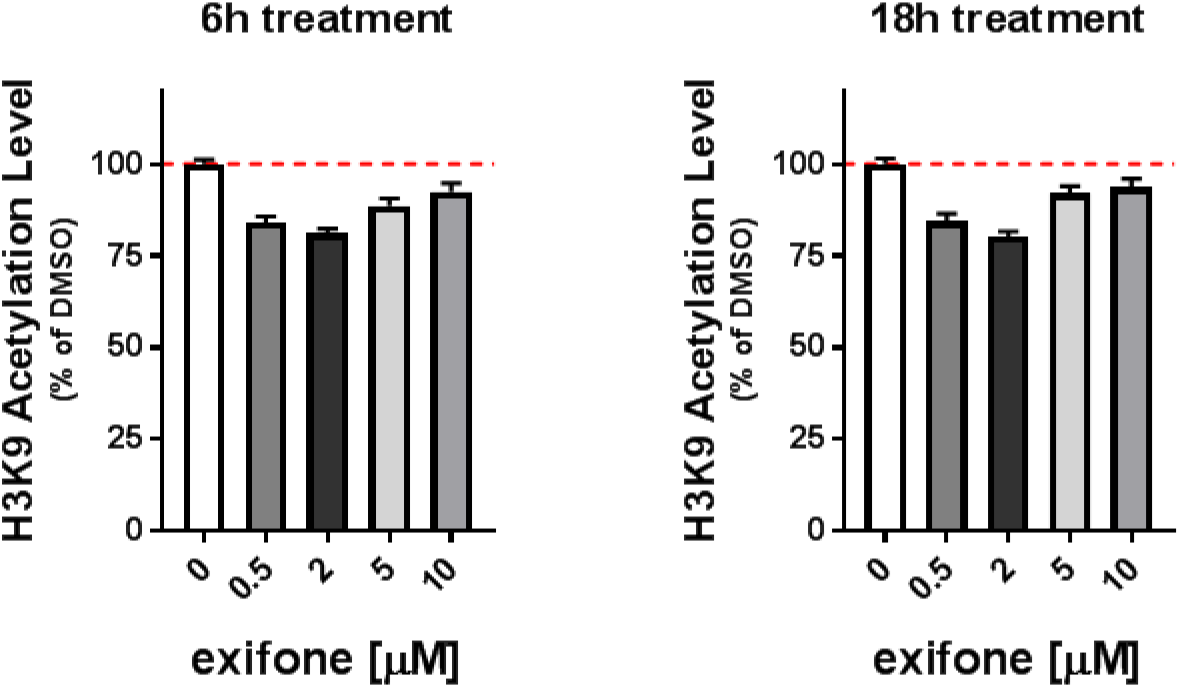
Exifone as an HDAC1 activator increases histone deacetylation of the H3K9Ac in human neural progenitor cells (NPCs). NPCs were treated with various concentrations of exifone for 6h or 18h, followed by fixation and immunostaining for H3K9Ac in nuclei. Quantitation of the H3K9Ac intensity levels indicates the activation of HDAC(s) in NPCs was most effectively by exifone at 0.5 μM and 2 μM. The amount of decrease of H3K9Ac levels became smaller with exifone at 5 or 10 μM, potentially indicating the loss of specificity of HDAC1 activation by exifone at those concentrations.

## References

1. Kim, D.; Frank, C. L.; Dobbin, M. M.; Tsunemoto, R. K.; Tu, W.; Peng, P. L.; Guan, J. S.; Lee, B. H.; Moy, L. Y.; Giusti, P.; Broodie, N.; Mazitschek, R.; Delalle, I.; Haggarty, S. J.; Neve, R. L.; Lu, Y.; Tsai, L. H., Deregulation of HDAC1 by p25/Cdk5 in neurotoxicity. Neuron 2008, 60 (5), 803–17.

2. Pao, P.-C.; Patnaik, D.; Watson, L. A.; Gao, F.; Pan, L.; Wang, J.; Adaikkan, C.; Penney, J.; Cam, H. P.; Huang, W.-C.; Pantano, L.; Lee, A.; Nott, A.; Phan, T. X.; Gjoneska, E.; Elmsaouri, S.; Haggarty, S. J.; Tsai, L.-H., HDAC1 modulates OGG1-initiated oxidative DNA damage repair in the aging brain and Alzheimer’s disease. Nature communications 2020, 11 (1), 2484.

3. Wang, W. Y.; Pan, L.; Su, S. C.; Quinn, E. J.; Sasaki, M.; Jimenez, J. C.; Mackenzie, I. R.; Huang, E. J.; Tsai, L. H., Interaction of FUS and HDAC1 regulates DNA damage response and repair in neurons. Nature neuroscience 2013, 16 (10), 1383–91.

4. Tsai, L.-H.; Pan, L.; Haggarty, S. J.; Patnaik, D. Activators of class I histone deacetylases (HDACs) and uses thereof. United States Patent 10,167,277, Jan 1, 2019.

5. Gazave J. M.; Rancurel A; Grenier., G. Certain biphenyl derivatives used to treat disorders caused by increased capillary permeability. Published March 29, 1977, US Patent Application 4,015,017A.

6. Piette F, S. S., Favre-Berrone M, Rainsard C, Poulain V, Rosen P, LaPeyre C, Berthaux P., Effets de I’exifone sur les troubles mnesiques, le comportement et al dépendance de malades atteints de démence sénile de type Alzheimer. La Revue de Gériatrie 1986, 11, 375–378.

7. Allain, H.; Denmat, J.; Bentue-Ferrer, D.; Milon, D.; Pignol, P.; Reymann, J. M.; Pape, D.; Sabouraud, O.; Van den Driessche, J., Randomized, double-blind trial of exifone versus cognitive problems in Parkinson’s disease. Fundamental & clinical pharmacology 1988, 2 (1), 1–12.

8. Porsolt, R. D.; Lenegre, A.; Avril, I.; Steru, L.; Doumont, G., The effects of exifone, a new agent for senile memory disorder, on two models of memory in the mouse. Pharmacology, biochemistry, and behavior 1987, 27 (2), 253–6.

9. Porsolt, R. D.; Lenegre, A.; Avril, I.; Lancrenon, S.; Steru, L.; Doumont, G., Psychopharmacological profile of the new cognition enhancing agent exifone in the mouse. Arzneimittel-Forschung 1987, 37 (4), 388–93.

10. Winter, R. W.; Cornell, K. A.; Johnson, L. L.; Ignatushchenko, M.; Hinrichs, D. J.; Riscoe, M. K., Potentiation of the antimalarial agent rufigallol. Antimicrobial agents and chemotherapy 1996, 40 (6), 1408–11.

11. Chichmanian, R. M.; Mignot, G.; Brucker, F.; Greck, T.; Spreux, A., [Exifone: 4 cases of hepatitis]. Gastroenterologie clinique et biologique 1989, 13 (4), 428–9.

12. Pariente, E. A.; Kapfer, J., [Hepatitis caused by exifone]. Gastroenterologie clinique et biologique 1989, 13 (4), 426.

13. Larrey, D., [Exifone, a new hepatotoxic drug?]. Gastroenterologie clinique et biologique 1989, 13 (4), 333–4.

14. Grange, J. D.; Biour, M.; Merrouche, Y.; Roland, J.; Bodin, F., [Hepatitis caused by exifone]. Gastroenterologie clinique et biologique 1989, 13 (4), 427–8.

15. Bentue-Ferrer, D.; Philouze, V.; Pape, D.; Reymann, J. M.; Allain, H.; Van den Driessche, J., Comparative evaluation of scavenger properties of exifone, piracetam and vinburnine. Fundamental & clinical pharmacology 1989, 3 (4), 323–8.

16. Largeron, M.; Lockhart, B.; Pfeiffer, B.; Fleury, M. B., Synthesis and in vitro evaluation of new 8-amino-1,4-benzoxazine derivatives as neuroprotective antioxidants. Journal of medicinal chemistry 1999, 42 (24), 5043–52.

17. Porsolt, R. D.; Lenegre, A.; Avril, I.; Doumont, G., Antagonism by exifone, a new cognitive enhancing agent, of the amnesias induced by four benzodiazepines in mice. Psychopharmacology 1988, 95 (3), 291–7.

18. Hertel, C.; Terzi, E.; Hauser, N.; Jakob-Rotne, R.; Seelig, J.; Kemp, J. A., Inhibition of the electrostatic interaction between beta-amyloid peptide and membranes prevents beta-amyloid-induced toxicity. Proceedings of the National Academy of Sciences of the United States of America 1997, 94 (17), 9412–6.

19. Blum, C. A.; Ellis, J. L.; Loh, C.; Ng, P. Y.; Perni, R. B.; Stein, R. L., SIRT1 modulation as a novel approach to the treatment of diseases of aging. Journal of medicinal chemistry 2011, 54 (2), 417–32.

20. Moniot, S.; Weyand, M.; Steegborn, C., Structures, substrates, and regulators of Mammalian sirtuins - opportunities and challenges for drug development. Frontiers in pharmacology 2012, 3, 16.

21. Segel, I. H., Enzyme kinetics: behavior and analysis of rapid equilibrium and steady state enzyme systems. 1975.

22. Dai, H.; Kustigian, L.; Carney, D.; Case, A.; Considine, T.; Hubbard, B. P.; Perni, R. B.; Riera, T. V.; Szczepankiewicz, B.; Vlasuk, G. P.; Stein, R. L., SIRT1 activation by small molecules: kinetic and biophysical evidence for direct interaction of enzyme and activator. J Biol Chem 2010, 285 (43), 32695–703.

23. Baptista, F. I.; Henriques, A. G.; Silva, A. M.; Wiltfang, J.; da Cruz e Silva, O. A., Flavonoids as therapeutic compounds targeting key proteins involved in Alzheimer’s disease. ACS chemical neuroscience 2014, 5 (2), 83–92.

24. Fass, D. M.; Shah, R.; Ghosh, B.; Hennig, K.; Norton, S.; Zhao, W. N.; Reis, S. A.; Klein, P. S.; Mazitschek, R.; Maglathlin, R. L.; Lewis, T. A.; Haggarty, S. J., Effect of Inhibiting Histone Deacetylase with Short-Chain Carboxylic Acids and Their Hydroxamic Acid Analogs on Vertebrate Development and Neuronal Chromatin. ACS Med Chem Lett 2010, 2 (1), 39–42.

25. Ghosh, B.; Zhao, W. N.; Reis, S. A.; Patnaik, D.; Fass, D. M.; Tsai, L. H.; Mazitschek, R.; Haggarty, S. J., Dissecting structure-activity-relationships of crebinostat: Brain penetrant HDAC inhibitors for neuroepigenetic regulation. Bioorg Med Chem Lett 2016, 26 (4), 1265–1271.

26. Zhao, W. N.; Ghosh, B.; Tyler, M.; Lalonde, J.; Joseph, N. F.; Kosaric, N.; Fass, D. M.; Tsai, L. H.; Mazitschek, R.; Haggarty, S. J., Class I Histone Deacetylase Inhibition by Tianeptinaline Modulates Neuroplasticity and Enhances Memory. ACS chemical neuroscience 2018, 9 (9), 2262–2273.

27. Silva, M. C.; Haggarty, S. J., Human pluripotent stem cell-derived models and drug screening in CNS precision medicine. Annals of the New York Academy of Sciences 2019.

28. Silva, M. C.; Cheng, C.; Mair, W.; Almeida, S.; Fong, H.; Biswas, M. H. U.; Zhang, Z.; Huang, Y.; Temple, S.; Coppola, G.; Geschwind, D. H.; Karydas, A.; Miller, B. L.; Kosik, K. S.; Gao, F. B.; Steen, J. A.; Haggarty, S. J., Human iPSC-Derived Neuronal Model of Tau-A152T Frontotemporal Dementia Reveals Tau-Mediated Mechanisms of Neuronal Vulnerability. Stem cell reports 2016, 7 (3), 325–340.

29. Vajda, F. J., Neuroprotection and neurodegenerative disease. Journal of clinical neuroscience : official journal of the Neurosurgical Society of Australasia 2002, 9 (1), 4–8.

30. Vogelauer, M.; Krall, A. S.; McBrian, M. A.; Li, J. Y.; Kurdistani, S. K., Stimulation of histone deacetylase activity by metabolites of intermediary metabolism. J Biol Chem 2012, 287 (38), 32006–16.

31. Singh, R. K.; Cho, K.; Padi, S. K.; Yu, J.; Haldar, M.; Mandal, T.; Yan, C.; Cook, G.; Guo, B.; Mallik, S.; Srivastava, D. K., Mechanism of N-Acylthiourea-mediated activation of human histone deacetylase 8 (HDAC8) at molecular and cellular levels. J Biol Chem 2015, 290 (10), 6607–19.

32. Tsai, L.-H.; Haggarty, S.; Patnaik, D.; Ping-Chieh, P., Compositions of polyhydroxylated benzophenones and methods of treatment of neurodegenerative disorders. US Patent App. 16/010,030: 2018.

33. Harrison, I. F.; Dexter, D. T., Epigenetic targeting of histone deacetylase: therapeutic potential in Parkinson’s disease? Pharmacology & therapeutics 2013, 140 (1), 34–52.

34. Beckers, T.; Burkhardt, C.; Wieland, H.; Gimmnich, P.; Ciossek, T.; Maier, T.; Sanders, K., Distinct pharmacological properties of second generation HDAC inhibitors with the benzamide or hydroxamate head group. International journal of cancer 2007, 121 (5), 1138–48.

35. Haggarty, S. J.; Tsai, L. H., Probing the role of HDACs and mechanisms of chromatin-mediated neuroplasticity. Neurobiology of learning and memory 2011, 96 (1), 41–52.

36. Bardai, F. H.; Price, V.; Zaayman, M.; Wang, L.; D’Mello, S. R., Histone deacetylase-1 (HDAC1) is a molecular switch between neuronal survival and death. J Biol Chem 2012, 287 (42), 35444–53.

37. Watson, P. J.; Millard, C. J.; Riley, A. M.; Robertson, N. S.; Wright, L. C.; Godage, H. Y.; Cowley, S. M.; Jamieson, A. G.; Potter, B. V.; Schwabe, J. W., Insights into the activation mechanism of class I HDAC complexes by inositol phosphates. Nature communications 2016, 7, 11262.

38. Decroos, C.; Christianson, N. H.; Gullett, L. E.; Bowman, C. M.; Christianson, K. E.; Deardorff, M. A.; Christianson, D. W., Biochemical and structural characterization of HDAC8 mutants associated with Cornelia de Lange syndrome spectrum disorders. Biochemistry 2015, 54 (42), 6501–13.

39. Deardorff, M. A.; Porter, N. J.; Christianson, D. W., Structural aspects of HDAC8 mechanism and dysfunction in Cornelia de Lange syndrome spectrum disorders. Protein science : a publication of the Protein Society 2016, 25 (11), 1965–1976.

40. Sperling, R. A.; Jack, C. R., Jr.; Aisen, P. S., Testing the right target and right drug at the right stage. Science translational medicine 2011, 3 (111), 111cm33.

41. Ryan, L.; Hay, M.; Huentelman, M. J.; Duarte, A.; Rundek, T.; Levin, B.; Soldan, A.; Pettigrew, C.; Mehl, M. R.; Barnes, C. A., Precision Aging: Applying Precision Medicine to the Field of Cognitive Aging. Frontiers in aging neuroscience 2019, 11, 128.

42. Boxer, A. L.; Gold, M.; Huey, E.; Hu, W. T.; Rosen, H.; Kramer, J.; Gao, F. B.; Burton, E. A.; Chow, T.; Kao, A.; Leavitt, B. R.; Lamb, B.; Grether, M.; Knopman, D.; Cairns, N. J.; Mackenzie, I. R.; Mitic, L.; Roberson, E. D.; Van Kammen, D.; Cantillon, M.; Zahs, K.; Jackson, G.; Salloway, S.; Morris, J.; Tong, G.; Feldman, H.; Fillit, H.; Dickinson, S.; Khachaturian, Z. S.; Sutherland, M.; Abushakra, S.; Lewcock, J.; Farese, R.; Kenet, R. O.; Laferla, F.; Perrin, S.; Whitaker, S.; Honig, L.; Mesulam, M. M.; Boeve, B.; Grossman, M.; Miller, B. L.; Cummings, J. L., The advantages of frontotemporal degeneration drug development (part 2 of frontotemporal degeneration: the next therapeutic frontier). Alzheimer’s & dementia : the journal of the Alzheimer’s Association 2013, 9 (2), 189–98.

43. Espay, A. J.; Kalia, L. V.; Gan-Or, Z.; Williams-Gray, C. H.; Bedard, P. L.; Rowe, S. M.; Morgante, F.; Fasano, A.; Stecher, B.; Kauffman, M. A.; Farrer, M. J.; Coffey, C. S.; Schwarzschild, M. A.; Sherer, T.; Postuma, R. B.; Strafella, A. P.; Singleton, A. B.; Barker, R. A.; Kieburtz, K.; Olanow, C. W.; Lozano, A.; Kordower, J. H.; Cedarbaum, J. M.; Brundin, P.; Standaert, D. G.; Lang, A. E., Disease modification and biomarker development in Parkinson disease: Revision or reconstruction? Neurology 2020, 94 (11), 481–494.

44. Ba, X.; Boldogh, I., 8-Oxoguanine DNA glycosylase 1: Beyond repair of the oxidatively modified base lesions. Redox biology 2017, 14, 669–678.

45. Rye, P. T.; Frick, L. E.; Ozbal, C. C.; Lamarr, W. A., Advances in label-free screening approaches for studying sirtuin-mediated deacetylation. J Biomol Screen 2011, 16 (10), 1217–26.

46. Rye, P. T.; Frick, L. E.; Ozbal, C. C.; Lamarr, W. A., Advances in label-free screening approaches for studying histone acetyltransferases. J Biomol Screen 2011, 16 (10), 1186–95.

47. Segel, I. H., Enzyme Kinetics: Behavior and Analysis of Rapid Equilibrium and Steady-state Enzyme Systems. John Wiley and Sons, Inc.: New York, 1975.

48. Shah, N. B.; Duncan, T. M., Bio-layer interferometry for measuring kinetics of protein-protein interactions and allosteric ligand effects. Journal of visualized experiments : JoVE 2014, (84), e51383.

